# Monitoring iron-sulfur cluster occupancy across the *E. coli* proteome using chemoproteomics

**DOI:** 10.1101/2021.04.01.438105

**Authors:** Daniel W. Bak, Eranthie Weerapana

## Abstract

Iron-sulfur (Fe-S) clusters are ubiquitous metallocofactors found across diverse protein families, where they perform myriad functions including redox chemistry, radical generation, and gene regulation. Monitoring Fe-S cluster occupancy in protein targets directly within native biological systems has been challenging. Commonly utilized spectroscopic methods to detect Fe-S clusters require purification of proteins prior to analysis. Global iron incorporation into the proteome can be monitored using radiolabeled iron, but limitations include the low resolution afforded by gel-based autoradiography. Here, we report the development of a mass spectrometry-based strategy to assess Fe-S cluster binding in a native proteome. This chemoproteomic strategy relies on monitoring changes in the reactivity of Fe-S cluster cysteine ligands upon disruption of Fe-S cluster incorporation. Application to *E. coli* cells cultured under iron-depleted conditions enabled monitoring of disruptions to Fe-S cluster incorporation broadly across the *E. coli* Fe-S proteome. Evaluation of *E. coli* deletion strains of three scaffold proteins within the Isc Fe-S biogenesis pathway enabled the identification of Fe-S clients that are reliant on each individual scaffold protein for proper cluster installation. Lastly, cysteine-reactivity changes characteristic of Fe-S ligands were used to identify previously unannotated Fe-S proteins, including the tRNA hydroxylase, TrhP, and a member of a family of membrane transporter ATPase subunits, DppD. In summary, the chemoproteomic strategy described herein provides a powerful tool to report on Fe-S cluster incorporation directly within a native proteome, to interrogate the role of scaffold and accessory proteins within Fe-S biogenesis pathways, and to identify previously uncharacterized Fe-S proteins.

---

Iron-sulfur (Fe-S) proteins harbor one or more Fe-S clusters, which are inorganic metallocofactors composed of iron and sulfide. Fe-S clusters are widely distributed across all species as one of the most evolutionarily ancient redox-active cofactors.^1^ Fe-S proteins are central components of essential cellular pathways, including: (1) electron-transfer reactions within respiratory chains;^2^ (2) radical generation for cofactor synthesis and tRNA/rRNA modification;^3-5^ (3) transcriptional regulation and DNA repair;^6-8^ and, (4) enzymatic generation of primary metabolites, such as aconitate and fumarate.^9,10^ In *E. coli* there are currently 144 annotated Fe-S proteins (**Supplemental Figure 1, Supplemental Table 1**), representing more than 3% of the *E. coli* proteome. This number is likely an underrepresentation as new Fe-S proteins are discovered every year.^11^ Fe-S clusters are typified by their ligation to protein cysteine residues, yet cysteine motifs and ligation patterns vary widely by protein family, making bioinformatic predictions of new Fe-S cluster binding sites particularly challenging.^12^ Even within the AdoMet radical enzyme family of Fe-S proteins, which typically contain a well-conserved CxxxCxxC ligation motif,^13,14^ atypical cysteine ligation patterns have recently been identified.^12,15^ As a result, the *de novo* identification of Fe-S proteins remains a challenge.

The biogenesis of Fe-S clusters is tightly regulated due to the high reactivity of the iron and sulfide species that comprise these clusters. The damaging toxicity of the cluster components has necessitated the evolution of intricate and tightly regulated pathways for the protected biosynthesis and delivery of these cofactors.^16-18^ For example, Fe-S cluster biogenesis in *E. coli* is carried out by the house-keeping Isc pathway and the stress responsive Suf pathway. These pathways shuttle Fe-S clusters among several scaffold proteins before ultimate transfer to client Fe-S proteins. Client proteins are either newly translated polypeptides, or proteins that have lost their Fe-S cluster due to oxidative damage,^19^ cannibalism of the cluster as a source of sulfur,^20,21^ or iron-limitation.^22^ Elucidating the exact role of each component within the Fe-S biogenesis pathway, and identifying the client proteins served by each scaffold protein, is complicated by the instability of Fe-S proteins and the inability to directly monitor Fe-S cluster occupancy within a native proteome. Currently, the most commonly used methods to examine cluster synthesis and delivery in native systems involve the use of radioactive isotopes of iron and sulfur, which is technically challenging, has limited sensitivity, and does not inform on the exact site of cluster binding within a protein.^23,24^ Alternatively, some aspects of Fe-S cluster biogenesis, delivery, and damage have been recapitulated with *in vitro* assays and purified Fe-S proteins,^25,26^ but it remains extremely challenging to confirm the physiological relevance of these *in vitro* studies.

Strategies to monitor Fe-S cluster binding directly in native systems would complement the existing methods to study Fe-S proteins. Such a platform will enable characterization of the function of Fe-S biogenesis components, and the identification of unannotated Fe-S client proteins. Here, we leverage a chemoproteomic technique to monitor changes in cysteine reactivity, termed isoTOP-ABPP,^27,28^ to assess the reactivity of cysteine ligands to Fe-S clusters from across the *E. coli* proteome. The isoTOP-ABPP platform applies a cysteine-reactive iodoacetamide-alkyne (IA-alkyne) probe to quantitatively report on differences in cysteine reactivity across two biological systems using LC-MS/MS. This strategy has been applied to identify cysteine residues that are targeted by covalent ligands,^29,30^ and reactive oxygen species.^31,32^ Oxidation and ligand binding result in a loss in cysteine labeling by the IA-alkyne probe. Similarly, it was demonstrated that binding to metal ions, specifically zinc, resulted in a loss in cysteine reactivity,^33^ alluding to the potential of applying isoTOP-ABPP to monitor metal binding to cysteine ligands on proteins. Here, we apply this cysteine-reactivity profiling strategy to inform on Fe-S cluster occupancy based on the distinct differences in the reactivity of Fe-S cysteine ligands in apo- versus holo-proteins. We demonstrate that isoTOP-ABPP is able to identify perturbations in the reactivity of Fe-S cysteine ligands on *E. coli* Fe-S proteins in response to iron-depletion, or deletion of components of the Isc pathway. We determine the subset of Fe-S cluster proteins affected by deletion of various scaffold proteins within the Isc pathway, and show that deletion of individual scaffold proteins have unique functional consequences on the global Fe-S proteome. Lastly, we identify cysteine residues that show reactivity signatures that align with known Fe-S cluster cysteine ligands, and determine two Fe-S proteins from *E. coli* that have not previously been characterized.

## Application of isoTOP-ABPP to monitor Fe-S cluster binding

We hypothesized that cysteine residues directly ligated to an Fe-S cluster on a holo-protein would show minimal reaction with IA-alkyne, whereas those same cysteines within the apo-protein should now be significantly more accessible to IA-alkyne labeling (**Figure 1A**). To confirm that there is a measurable difference in IA-alkyne labeling of Fe-S cysteine ligands in apo- versus holo-proteins, we focused on the [4Fe-4S] cluster-containing protein aconitase (AcnA). AcnA was overexpressed and purified from *E. coli* (BL21) cells grown under standard (iron-replete) and iron-depleted conditions. Only AcnA grown under iron-replete conditions displayed a typical UV-visible absorbance spectrum associated with the presence of a [4Fe-4S] cluster (**Figure 1B**). The purified proteins were treated with IA-alkyne, conjugated to a fluorophore using copper-catalyzed azide-alkyne cycloaddition (CuAAC), and subsequently visualized by in-gel fluorescence. IA-alkyne labeling of AcnA was observed only when purified from the iron-depleted growth condition, while negligible labeling was observed under iron-replete conditions (**Figure 1C**). These studies confirmed that the cysteine ligands for the Fe-S cluster on AcnA displayed significantly enhanced reactivity within the apo-protein relative to the holo-protein.

**Fig. 1.**
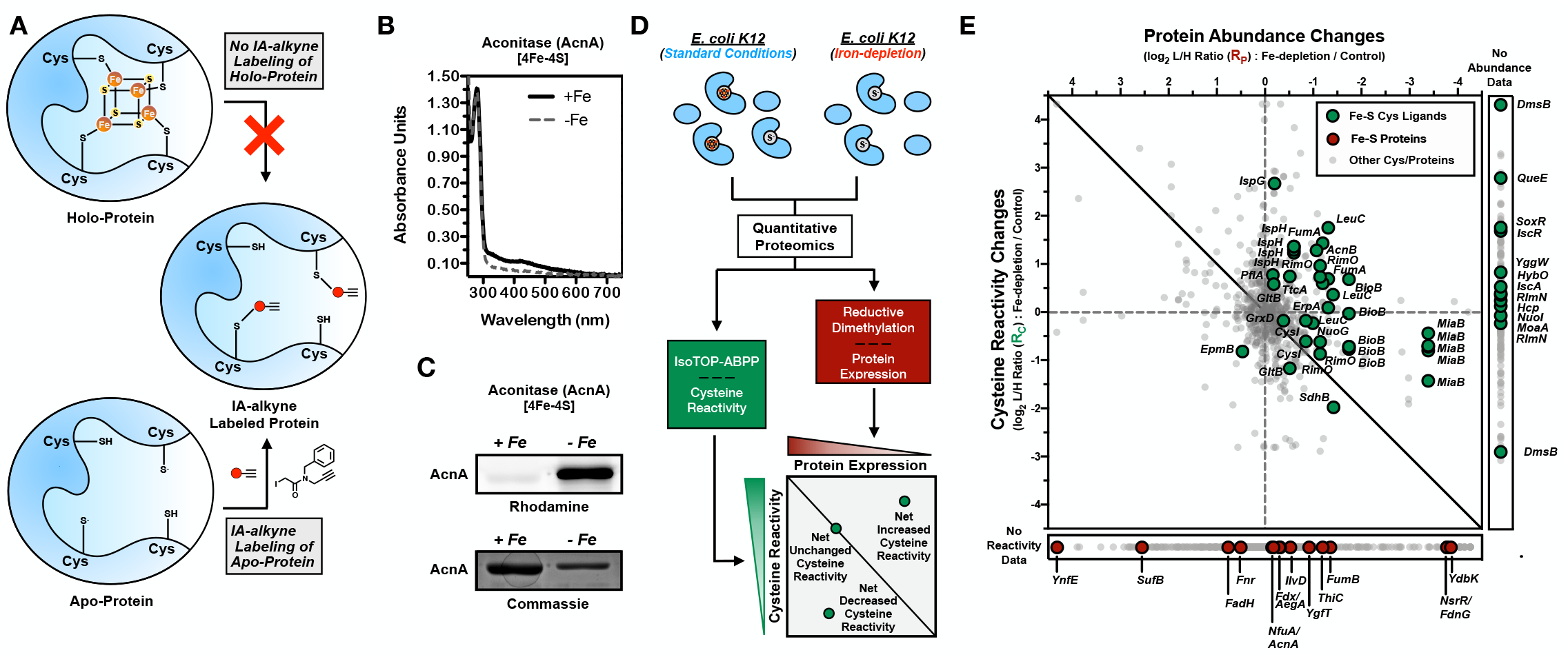
A two-dimensional chemoproteomic platform to monitor Fe-S cluster binding. **a**, Labeling of apo- and holo-proteins with a cysteine-reactive IA-alkyne probe. **b**, UV-visible absorbance spectrum of recombinant aconitase (AcnA) purified from iron-depleted (--) or iron-replete (—) media. **c**, Rhodamine and coomassie-blue gels of IA-alkyne labeled AcnA purified from iron-depleted (-Fe) or iron-replete (+Fe) media. **d**, Two-dimensional proteomic platform for monitoring net changes in cysteine reactivity across a bacterial proteome during iron-limitation. Quantitative chemoproteomic methods are applied to measure cysteine labeling (isoTOP-ABPP) and protein abundance (ReDiMe) changes upon iron-depletion. Cysteine residues with ratios above the expected linear correlation line are considered to have a net increase in cysteine reactivity, those on the line no net change, and those below the line a net decrease in cysteine reactivity. **e**, Two-dimensional proteomic dataset for the *E. coli* proteome grown under iron-depletion. All quantified cysteine residues are plotted in the main graph (annotated Fe-S cluster cysteine ligands - green circles, non-annotated cysteine residues - light gray small circles). Inset to right: cysteine residues with no protein abundance data (annotated Fe-S cluster cysteine ligands - green circles, non-annotated cysteine residues - light gray small circles). Inset below: proteins with no cysteine reactivity data (annotated Fe-S protein - red circles, non-annotated proteins - light gray small).

Given that AcnA displayed the expected increase in cysteine reactivity in iron-depleted conditions, we proceeded to determine if this trend was maintained across the *E. coli* Fe-S proteome using the isoTOP-ABPP platform. In most prior applications of isoTOP-ABPP, cells were treated with ligand or oxidant for short periods of time, where changes in protein expression were unlikely to be significant. In contrast, it is well known that *E. coli* cells respond to iron availability by activating a transcriptional program geared to increase intracellular iron levels.^34^ Therefore, any changes in IA-alkyne labeling would reflect on changes in both cysteine reactivity and protein abundance. To account for the protein abundance changes, and enable the quantification of net changes in cysteine reactivity, we developed a two-dimensional chemoproteomic strategy. This two-dimensional analysis quantitatively monitors changes in protein abundance using reductive dimethylation (ReDiMe) in the first dimension, and changes in IA-alkyne labeling of cysteines using isoTOP-ABPP in the second dimension. Cysteines showing no net change in reactivity should track linearly with protein abundance, and lie along the diagonal linear correlation line in the two-dimensional plot (**Figure 1D**). Cysteines displaying net increases in reactivity are located above the linear correlation line, and the further the distance from the diagonal, the greater the net change in cysteine reactivity. Dysregulation of Fe-S cluster incorporation would lead to net increases in cysteine reactivity for Fe-S ligating cysteines, where the magnitude of the net increase is indicative of the relative abundance of apo- versus holo-protein.

We first applied our two-dimensional chemoproteomic analysis to monitor net changes in cysteine reactivity for cysteine ligands on Fe-S proteins from *E. coli* cultured under iron-replete and iron-depleted conditions. *E. coli* (K12) cells were grown in minimal media with standard (iron-replete) or 100-fold decreased (iron-depleted) levels of iron. During exponential growth (OD_600_ ∼ 0.6), this resulted in ∼3-fold decrease in intracellular iron levels as measured by inductively coupled plasma-optical emission spectrometry (ICP-OES) (**Supplemental Figure 2A**). In contrast, intracellular zinc levels remained constant (**Supplemental Figure 2B**), confirming the specific depletion of iron over other metals. Cells were harvested during exponential growth phase and cell lysates were processed for both ReDiMe^35^ (**Supplemental Figure 2C**) and isoTOP-ABPP^27^ (**Supplemental Figure 2D**) analyses. In the ReDiMe dimension, proteins were subjected to trypsin digestion and isotopic labeling through reductive dimethylation with light (iron-depleted) and heavy (iron-replete) formaldehyde. The resulting protein log_2_ light: heavy (L/H) ratios, R_P_, reflect the relative protein abundance under the two conditions, where R_P_ > 0 indicates an increase in protein abundance under iron depletion. In the isoTOP-ABPP dimension, iron-depleted and iron-replete cell lysates were labeled with isotopically tagged variants of the IA-alkyne probe,^36^ IA-light and IA-heavy (**Supplemental Figure 2D**), respectively. IA-labeled proteins were conjugated to chemically cleavable biotin-azide using CuAAC, and the light and heavy-labeled samples were combined, enriched on streptavidin beads, and subjected to on-bead trypsin digestion. IA-labeled peptides were then released by sodium dithionite treatment and analyzed by LC/LC-MS/MS. The resulting cysteine log_2_ L/H ratios, R_C_, reflect the relative changes in cysteine labeling, where R_C_ > 0 indicates an increase in cysteine labeling in the iron-depleted sample.

In the ReDiMe analysis, the relative abundance changes in iron-depleted and iron-replete conditions were quantified for 1,422 proteins, with 479 displaying increased or decreased protein abundance (R_P_ > |0.5|) (**Supplemental Figure 2E, Supplemental Table 2**). As confirmation that these changes in protein abundance were the direct result of iron depletion, gene-ontology analysis of the 247 up-regulated proteins (R_P_ > 0.5) showed significant enrichment of processes involved in iron uptake, siderophore biosynthesis, and metal homeostasis (**Supplemental Figure 2F**). Importantly, 42 Fe-S proteins were identified, of which 24 were down-regulated (**Supplemental Figure 2G)**. This downregulation of Fe-S proteins under iron depletion has previously been noted to be a downstream consequence of the global iron-responsive transcription factor, Fur.^34^ This significant perturbation of Fe-S protein levels under iron depletion underscores the importance of accounting for protein abundance changes in the typical isoTOP-ABPP workflow.

In the isoTOP-ABPP analysis, differences in IA-alkyne labeling in iron-depleted and iron-replete *E. coli* proteomes were reported for 896 cysteines, where 651 had corresponding protein abundance information from the ReDiMe analysis **(Figure 1E, Supplemental Table 2)**. In total, 49 annotated Fe-S ligating cysteine residues were identified in the isoTOP-ABPP analysis, and of these, 29 had corresponding protein abundance changes determined by ReDiMe analysis (**Figure 1E**). The remaining 20 Fe-S ligating cysteines could not be corrected for protein abundance, since the corresponding proteins were not detected in the ReDiMe analysis. Similarly, 15 Fe-S proteins identified in the ReDiMe analysis did not generate cysteine-labeling data by isoTOP-ABPP. In general, Fe-S ligating cysteines displayed net increases in cysteine reactivity, compared to the entire dataset (**Figure 2A**). Additionally, to demonstrate that the observed net changes in cysteine reactivity were driven by Fe-S cluster occupancy, net cysteine reactivity changes for non Fe-S ligating cysteines on Fe-S proteins displayed no changes in net cysteine reactivity (**Figure 2A)**. In these instances, we can adapt the average R_C_ values for non-ligating cysteines in each protein as a proxy for protein abundance, and further increase our total coverage in our two-dimensional plot to 36 Fe-S ligating cysteines. Lastly, to demonstrate the quantitative accuracy of the analysis, a control sample of standard versus standard growth conditions showed no changes in protein abundance by ReDiMe, cysteine labeling by isoTOP-ABPP, and by extension no net changes in cysteine reactivity (**Supplemental Figure 2H, Supplemental Table 3**). Together, these analyses confirm that net changes in cysteine reactivity are a meaningful proxy for Fe-S occupancy, and the combined ReDiMe and isoTOP-ABPP analysis can reflect changes in Fe-S occupancy across the proteome.

**Fig. 2.**
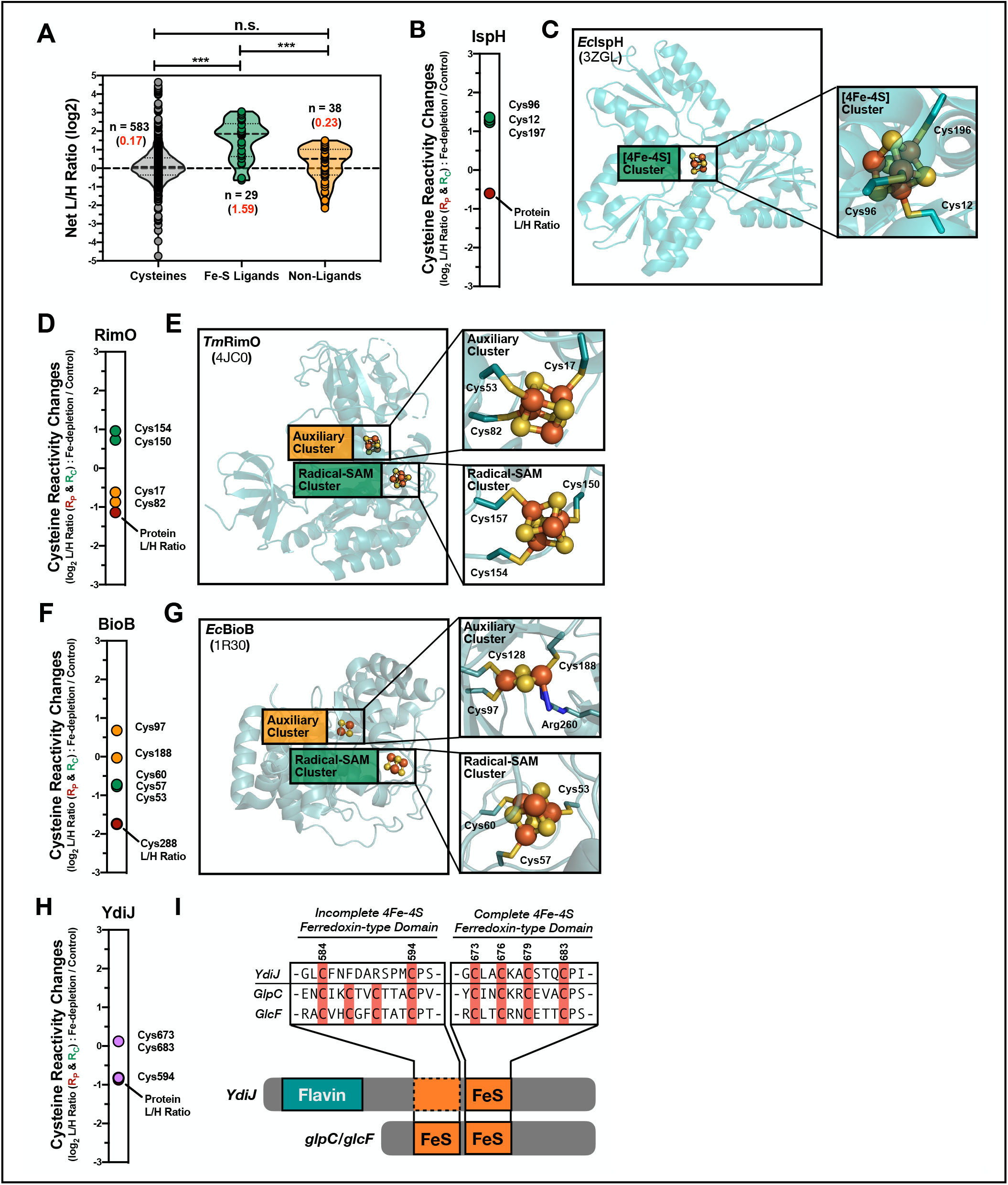
Monitoring reactivity changes for Fe-S ligating cysteines. **a**, Violin plot of Net L/H ratios (R) for Fe-S ligands (green), non-liganding cysteines from Fe-S proteins (yellow), and all other cysteine residues from non-Fe-S proteins (gray). The median R value is displayed as a dashed line in the violin plot, while the average R values (red) and number of unique values (black) are indicated for each group. Significance is calculated as *** p < 0.005, paired t-test (two-tailed), with n indicated in the figure. **b, d, f, h**, Rc values for Fe-S cluster ligating cysteines on Fe-S proteins; **b**, IspH, **d**, RimO **f**, BioB, and **h**, YdiJ. **c, e, g** Crystal structures of each Fe-S proteins with cluster and cysteine residues highlighted, with cluster and cysteine residues highlighted; **c**, *E. coli* IspH (Inset: [4Fe-4S] cluster – green), **e** RimO from *Thermotoga maritima* (Insets: AdoMet radical [4Fe-4S] cluster – green and auxiliary [4Fe-4S] cluster – orange), and **g**, *E. coli* BioB (Insets: AdoMet radical [4Fe-4S] cluster – green and auxiliary [4Fe-4S] cluster – orange). **i**, Alignment of YdiJ, with homologs, GlpC and GlcF. Two [4Fe-4S] ferredoxin-type domains are shown, one complete and one incomplete.

Detailed analyses of net changes in cysteine reactivity for individual proteins afforded some interesting insight into the properties of different Fe-S clusters. The 49 annotated Fe-S ligating cysteines we identified by isoTOP-ABPP were localized on 32 Fe-S clusters. We observed IA-alkyne labeling of multiple cysteines within a cluster for 12 (38%) of the Fe-S clusters identified. When multiple cysteines within a cluster were identified, each individual cysteine displayed a similar magnitude of reactivity change. For example, the 3 cysteine ligands of the 4-hydroxy-3-methylbut-2-enyl diphosphate reductase, IspH, Fe-S cluster, uniformly displayed R_C_ values of ∼1.2 (**Figure 2B,C**). For proteins that contain multiple Fe-S clusters, cysteines within each individual cluster showed differential net changes in reactivity. For example, the ribosomal protein S12 methylthiotransferase, RimO, contains two Fe-S clusters, an AdoMet radical [4Fe-4S] cluster, and an auxiliary [4Fe-4S] cluster. Cysteine ligands for the auxiliary cluster displayed a modest net increase in reactivity, while cysteine ligands for the AdoMet radical cluster displayed a ∼ 4-fold net increase in reactivity (**Figure 2D,E**). In contrast, for biotin synthase (BioB), the auxiliary [2Fe-2S] cluster displays a greater (∼4-fold) net increase in cysteine reactivity compared to the AdoMet radical cluster (∼3-fold) (**Figure 2F,G**). Therefore, the auxiliary cluster of BioB appears to be significantly less stable than the auxiliary cluster of RimO, relative to the AdoMet radical clusters on each protein. These observations are consistent with the reaction mechanisms of these two enzymes, where the auxiliary cluster of BioB is sacrificed as a source of sulfur during the course of the reaction cycle,^20,21^ while that of RimO is left intact.^37^ Lastly, this platform allows for the tentative assignment of ligation sites in Fe-S proteins that have yet to be fully characterized. For example, YdiJ was recently identified as a putative Fe-S protein through bioinformatics, and subsequently confirmed experimentally to bind a single Fe-S cluster (likely [4Fe-4S]).^12^ However, the exact cysteine ligands for the Fe-S cluster were not assigned. We quantified the net changes in reactivity for three cysteine residues on YdiJ (C594, C673, and C683). Of these, C673 and C683 displayed a significant net increase in cysteine reactivity, with C594 showing no net change in reactivity, suggesting that C673 and C683 are likely to be Fe-S ligating residues on YdiJ (**Figure 2H**). This observation is supported by the fact that C673 and C683 belong to a conserved ferredoxin-like motif (as compared to the *E. coli* homologs, GlcF and GplC), while C594 is part of a partial ferredoxin-like motif that appears incomplete in this protein (**Figure 2I)**. Together, our data demonstrate the ability of our two-dimensional chemoproteomic platform to: (1) report on Fe-S cluster occupancy across multiple Fe-S proteins within the *E. coli* proteome; (2) compare the relative occupancy levels of multiple Fe-S clusters within a single protein; and, (3) assign Fe-S cluster ligating cysteine residues on Fe-S proteins where the exact site of cluster binding is undetermined.

### Interrogating Fe-S cluster transfer and delivery

Upon confirming the ability of our two-dimensional analyses to report on changes in Fe-S cluster binding upon iron-depletion, we sought to apply our platform to *E. coli* proteomes where Fe-S biogenesis was disrupted in a more targeted manner without global perturbations of cellular iron. We therefore turned to *E. coli* genetic deletion strains of the various scaffold proteins of the Isc Fe-S cluster biogenesis pathway (**Figure 3A**), including: (1) IscU, the initial scaffold protein that is primarily responsible for [2Fe-2S] cluster synthesis; (2) IscA, an A-type scaffold that synthesizes and delivers [4Fe-4S] clusters; and, (3) NfuA, an alternative scaffold protein involved in targeted cluster delivery or repair of damaged/lost Fe-S clusters.^16^ We applied our two-dimensional chemoproteomic analysis to these deletion strains, where we compared *ΔiscU* (**Figure 3B, Supplemental Table 4**), *ΔiscA* (**Figure 3C, Supplemental Table 5**), and *ΔnfuA* (**Figure 3D, Supplemental Table 6**), to wild-type *E. coli* (K12) cells. We identified 38, 24, and 31 Fe-S ligating cysteines in the *ΔiscU, Δisc*A, and *ΔnfuA* isoTOP-ABPP analyses, respectively. As observed with iron-depletion, the majority of Fe-S ligating cysteine residues consistently displayed net increases in cysteine reactivity, albeit with varying magnitude depending on the scaffold protein that is deleted.

**Fig. 3.**
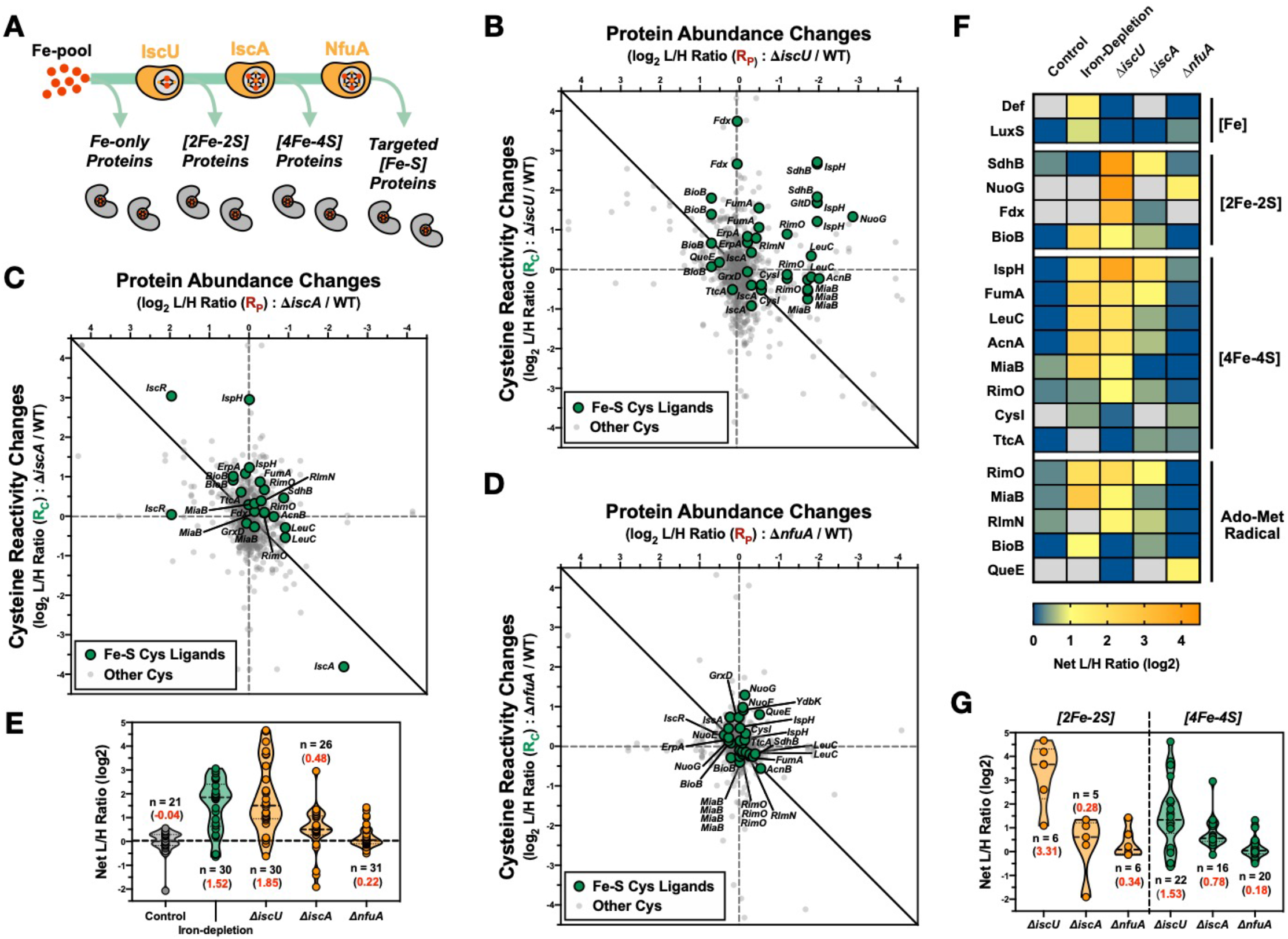
Interrogating the role of scaffold proteins within the Isc Fe-S biogenesis pathway. **a**, Simplified scheme of Isc-mediated Fe-S biogenesis. **b-d**, Two-dimensional proteomic datasets for the *E. coli ΔiscU* (**b**), *ΔiscA* (**c**), and *ΔnfuA* (**d**) strains (annotated Fe-S cluster cysteine ligands - green circles, non-annotated cysteine residues - light gray small circles). **e**, Violin plot of net changes in cysteine reactivity for all Fe-S cluster cysteine ligands from control (gray), iron-depletion (green), and Isc deletion strains (orange). Number of values (n) and average corrected L/H ratio (red) are indicated for each group.**f**, Heat map of Fe-S clusters (and iron-binding sites) across all experimental growth conditions and cell strains. Each cell represents the average net change in cysteine reactivity from all quantified cysteines for a given Fe-S cluster. **g**, Violin plot of net changes in cysteine reactivity for all [2Fe-2S] (orange) and [4Fe-4S] (green) cluster cysteine ligands from the *ΔiscU, ΔiscA. and ΔnfuA* datasets. Number of values (n) and average corrected L/H ratio (red) are indicated for each group.

We initially compared the four datasets generated (iron-depletion, *ΔiscU, ΔiscA, ΔnfuA*) (**Figure 3E,F)**, to understand how external (iron-depletion) and internal (*ΔiscU, ΔiscA, ΔnfuA*) disruptions to Fe-S biogenesis differentially affect the Fe-S proteome. One key difference between iron-depletion and the genetic disruption of Fe-S biogenesis, is the cysteine-reactivity change observed for two non-heme iron-only proteins, peptide deformylase (Def) and S-ribosylhomocysteine lyase (LuxS), which each contain a single cysteine residue ligated to the iron center.^38,39^ Both of these cysteine residues displayed net increases in reactivity only under iron-depletion conditions, and not in any of the *isc*-deletion strains. This is expected, as the Isc pathway is specific for the synthesis and delivery of Fe-S clusters, and is not implicated in iron delivery to non Fe-S proteins (**Figure 3A**). When comparing the *ΔiscU, ΔiscA*, and *ΔnfuA* datasets, it is clear that the *ΔiscU* strain has the strongest effect on the Fe-S proteome, with the *ΔiscA* and *ΔnfuA* strains displaying modest to minimal changes, respectively (**Figure 3E,F**). These observations are consistent with the known roles of these scaffold proteins within the pathway, where IscU is situated in the early phase of Fe-S biogenesis (**Figure 3A**), and therefore the *ΔiscU* strain is more likely to have large global perturbations to the Fe-S proteome. In contrast, IscA and NfuA are situated in the later phases of Fe-S biogenesis and are expected to have a less significant impact on the Fe-S proteome. Lastly, in the *ΔiscU* strain, the largest net changes in cysteine reactivity are observed for [2Fe-2S] clusters. In contrast, these same [2Fe-2S] clusters are minimally affected in the *ΔiscA* and *ΔnfuA* strains (**Figure 3G**). This is in agreement with the characterized organization of the pathway, where [2Fe-2S] clusters are generated on IscU, whereas [4Fe-4S] clusters are generated on IscA (**Figure 3A**). Therefore, all [2Fe-2S] cluster-containing client proteins should be directly dependent on the presence of IscU, but not IscA or NfuA.

We then sought to utilize our data to delineate specific pathways for cluster delivery to individual Fe-S client proteins. First, it is important to note that there are two Fe-S biogenesis pathways in *E. coli*; the Isc pathway, and the stress-inducible Suf pathway.^18^ Both Isc and Suf proteins are expressed from their respective operons, with the *suf* operon being induced by the Fe-S sensing transcription factor IscR^40^ (**Supplemental Figure 3A and B**). Perturbations in Fe-S biogenesis that lead to the formation of apo-IscR result in activation of the Suf system. Using the data from our ReDiMe analysis, we can determine the relative expression of components of the Isc (IscR, S, U and A) and the Suf (SufA, B, C, D, and S) pathways under the four conditions analyzed (iron-depletion, *ΔiscU, ΔiscA, ΔnfuA*). Notably, under conditions of iron-depletion, and in the *ΔiscU* strain, we observe significant Suf induction (**Supplemental Figures 3C**). In contrast, lower levels of Suf induction are observed in the *ΔiscA* and *ΔnfuA* strains, likely because IscU is still active and able to form the [2Fe-2S] cluster on IscR to suppress Suf activation (**Supplemental Figure 3B**). We were interested to determine if the Suf pathway is able to serve all, or only a subset, of Fe-S client proteins. Given the essentiality of IscU to the Isc pathway, we can consider the *ΔiscU* strain as defective in the Isc pathway, and only able to load Fe-S clients through the Suf pathway. In the *ΔiscU* strain, we observed a number of Fe-S clusters that were minimally disrupted by loss of the Isc pathway (**Figure 3B**). We therefore hypothesize that these clusters are targets of the Suf pathway. These putative Suf targets include Fe-S clusters of CysI, QueE, TtcA and BioB client proteins.

Similarly, we sought to assign client proteins for different scaffold components within the Isc pathway. Specifically, we focused on the two A-type scaffold proteins in the Isc pathway, IscA and NfuA, and directly compared the datasets for the *ΔiscA* and *ΔnfuA* strains (**Figure 3C and D**). In these strains, the Suf pathway is not activated, and therefore the only means of obtaining cluster is through components of the Isc pathway. Relative to iron-depletion conditions and the *ΔiscU* strain, fewer perturbations to the Fe-S proteome were identified in the *ΔiscA* strain (**Figure 3F**). This is indicative of the ability to circumvent IscA within the Fe-S biogenesis pathway, and utilize other scaffold proteins to form the cluster on the majority of client proteins. However, significant disruptions to Fe-S cluster formation were observed for IspH in particular (**Figure 3C**), suggesting that IspH is a privileged client for IscA over other scaffold proteins. The *ΔnfuA* strain showed minimal perturbations in Fe-S cluster formation across the Fe-S proteome (**Figure 3F**). The most significant perturbations in Fe-S cluster formation in the *ΔnfuA* strain were for NuoG, NuoF, QueE and YdbK (**Figure 3D**). Of these, NuoG and NuoF are subunits of the respiratory NADH dehydrogenase complex. A previous study reported decreased NADH dehydrogenase activity in the *ΔnfuA* strain,^41^ supporting our observation that NuoG and NuoF are likely privileged clients of the NfuA scaffold protein. NfuA is also responsible for the repair or delivery of the auxiliary [4Fe-4S] cluster of the Ado-Met radical enzyme lipoyl synthase (LipA).^42^ However, LipA was not identified in our data. Together, our data enable the identification of client proteins dependent on different A-type scaffolds, specifically showing that IspH is a client for IscA, and NuoG and NuoF are clients for NfuA.

### Mapping the coverage and reactivity of *E. coli* Fe-S cysteine ligands

To determine the overall coverage of the *E. coli* Fe-S proteome accessible through our chemoproteomic platform, we combined the data from our four analyses (iron-depletion, *ΔiscU, ΔiscA, ΔnfuA*). The *E. coli* Fe-S proteome is comprised of 144 members that can be divided into 6 functional categories of Fe-S cluster proteins (**Figure 4, Supplemental Figure 1**): (1) ferredoxin-like cluster containing proteins (cluster involved in electron transfer – fully coordinated); (2) AdoMet radical enzymes (cluster involved in radical generation – open coordination site occupied by S-adenosylmethionine); (3) dehydratases (cluster involved in non-redox catalysis – open coordination site for substrate binding); (4) DNA-binding proteins (cluster involved in transcriptional regulation, long range electron transfer or redox sensing); (5) Fe-S Scaffold proteins (cluster is formed and transferred to client Fe-S proteins); and, (6) miscellaneous Fe-S proteins. Across our four two-dimensional analyses, 94 of the 144 Fe-S proteins were identified in either the isoTOP-ABPP or the ReDiMe analyses. Fe-S clusters and proteins from all 6 functional categories were observed, with varying coverage across the 5 datasets (**Supplemental Figure 4A**). In general, ferredoxin-like clusters, often part of large respiratory complexes, were less well represented relative to other functional categories (**Supplemental Figure 4B**). This is likely due to the fact that these respiratory complexes are often only expressed under anaerobic conditions, or in the presence of very specific electron acceptors.^43^ Strong coverage is observed for Fe-S proteins in the aerobic respiratory pathway (**Supplemental Figure 4C**), in contrast to the Fe-S proteins from alternative respiratory pathways, such as those involved in hydrogen and formate metabolism (**Supplemental Figure 4D**). It is likely that coverage of the Fe-S proteome could be further improved by profiling cells grown under anaerobic conditions or on alternative nutrient sources. Another reason for reduced coverage of the ferredoxin-like clusters is likely due to these clusters being deeply imbedded into the protein core,^44^ with reduced accessibility to IA-alkyne. In contrast, catalytic and scaffold Fe-S clusters are often more surface exposed^45,46^, and thereby accessible to small-molecule substrates, and, by extension to IA-alkyne.

**Fig. 4.**
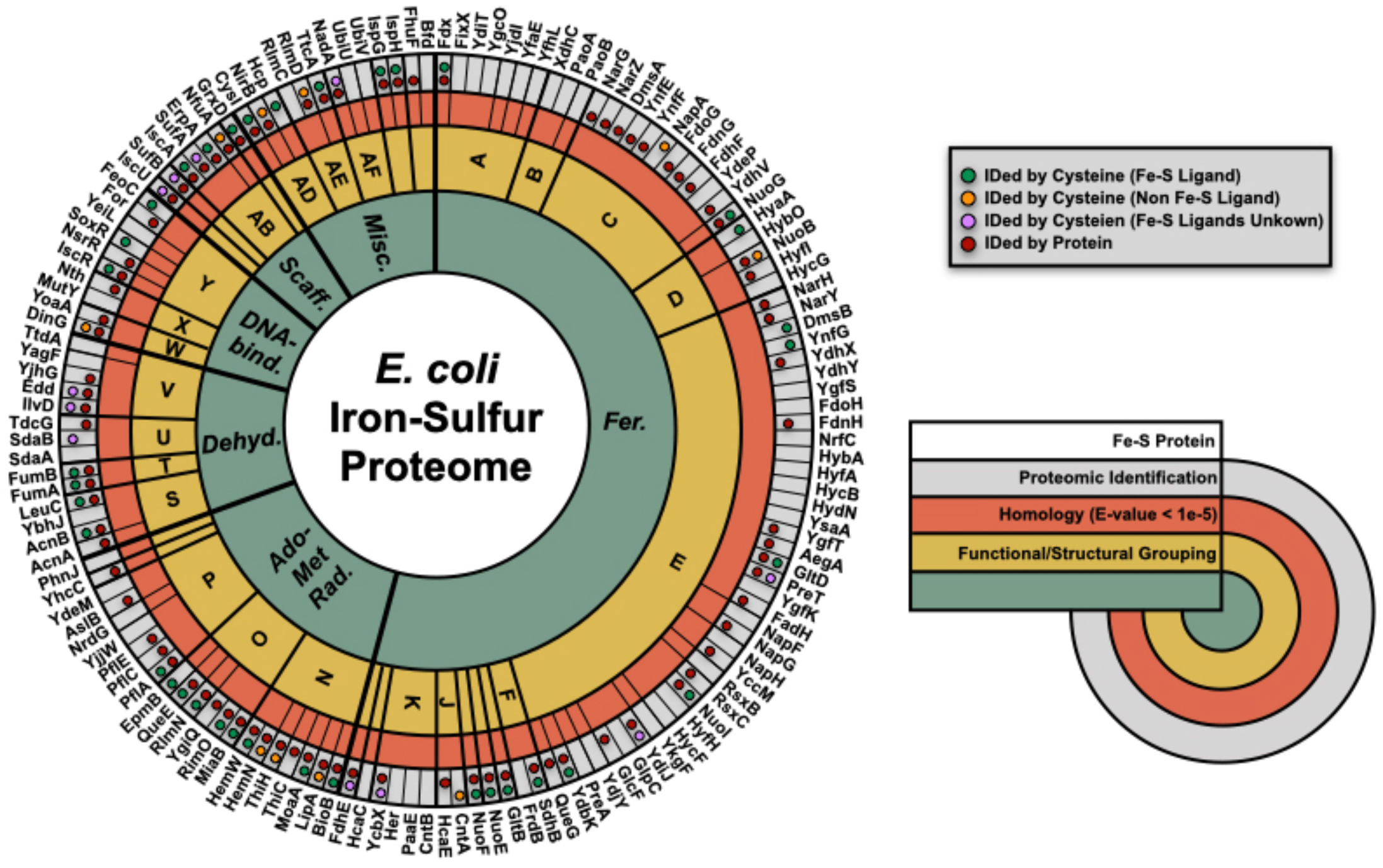
Mapping coverage of the *E. coli* Fe-S proteome. Wheel diagram of the 144-member annotated *E. coli* Fe-S Proteome. The *E. coli* Fe-S proteome is divided into 6 functional families (inner first ring – green), smaller sub-functional and structural groups (second ring – yellow), and homology-related (E-value < 1E-5) clusters (third ring – orange). Protein identification by the 2-dimensional proteomic platform is indicated (fourth outer ring – gray) for protein abundance (red circles) and cysteine reactivity (annotated Fe-S cluster cysteine ligands - green circles, non-Fe-S ligating cysteine residues – orange circles, cysteine residues, where Fe-S ligands are unknown for the protein – purple circle).

The fact that we identify a significant number of Fe-S ligating cysteines in our isoTOP-ABPP analysis, suggests that these cysteines are successfully competing with other cellular cysteines for the limited pool of IA-alkyne that is available for cysteine labeling. We were therefore interested in characterizing the inherent reactivity of an Fe-S ligating cysteine relative to other functional cysteines in the proteome. To determine the inherent reactivity of Fe-S ligating cysteines, we performed an isoTOP-ABPP analysis geared to rank IA-alkyne labeled cysteines by relative reactivity.^27^ Iron-depleted *E. coli* lysates were treated with either low (10 µM) or high (100 µM) concentration of IA-heavy and IA-light, respectively. Highly reactive cysteines are expected to saturate labeling at the low IA-alkyne concentration, generating L/H ratios of ∼1. Therefore, lower L/H ratio values indicate increased reactivity of a cysteine residue (**Supplemental Figure 5A**). This analysis revealed that while Fe-S ligating cysteine residues displayed a range of reactivity, many were found to be highly reactive with L/H ratios < 5 (**Supplemental Figure 5B, Supplemental Table 7**). In fact, cysteines involved in Fe-S ligation displayed increased reactivity relative to zinc-binding and disulfide-linked cysteines, and were only minimally less reactive than active-site cysteine nucleophiles (**Supplemental Figure 5C**). An interesting distinction was observed across Fe-S ligating cysteines, where Fe-S ligands in client proteins were more reactive than those in scaffold proteins involved in Fe-S cluster biogenesis (**Supplemental Figure 5D**). This difference in reactivity likely reflects on the stable versus transient chelation to the cluster that is characteristic of client and scaffold proteins, respectively. Together, our reactivity profiling studies demonstrate that Fe-S cluster ligands show high inherent reactivity when not bound to the Fe-S cluster, and are therefore likely susceptible to modification by reactive oxygen species and electrophiles.

### Identification of unannotated bacterial Fe-S proteins

We next explored the promise of our chemoproteomic strategy to identify previously uncharacterized Fe-S proteins. As stated previously, it is challenging to predict Fe-S cysteine ligands, as many Fe-S proteins have poorly conserved primary sequence motifs. Even for known Fe-S proteins, assigning the exact cysteine ligands for an Fe-S cluster is difficult without access to high-resolution structural data. The unbiased nature of our chemoproteomic platform allows us to monitor changes to cysteine reactivity across residues and proteins that are not otherwise predicted or anticipated to be involved in Fe-S cluster binding. To arrive at a list of putative uncharacterized iron-binding proteins, we filtered our data in the following manner. First, we identified a subset of 142 cysteines that displayed a greater than 3-fold net increase in reactivity in at least one of the four conditions (iron-depletion, *ΔiscU, ΔiscA, ΔnfuA*). Upon removing known Fe-S proteins, we then further filtered for proteins that contain at least three cysteine residues, a requirement for Fe-S cluster ligation, thereby reducing our target list to 58 cysteines on 46 proteins (**Supplemental Figure 6)**. Lastly, we generated a high-confidence list of 18 cysteines on 14 proteins, by requiring a net increase in reactivity in at least two of the four datasets acquired (**Figure 5A**). It is important to recognize that this list of proteins could contain cysteine residues that show changes in reactivity resulting from conformational changes driven by Fe-S occupancy within a complexing protein partner. For example, the cysteine residues on the NADH dehydrogenase subunit, NuoC, which does not bind one of the nine Fe-S clusters of the complex,^44^ may show reactivity changes resulting from the loss of other clusters within the complex.

**Fig. 5.**
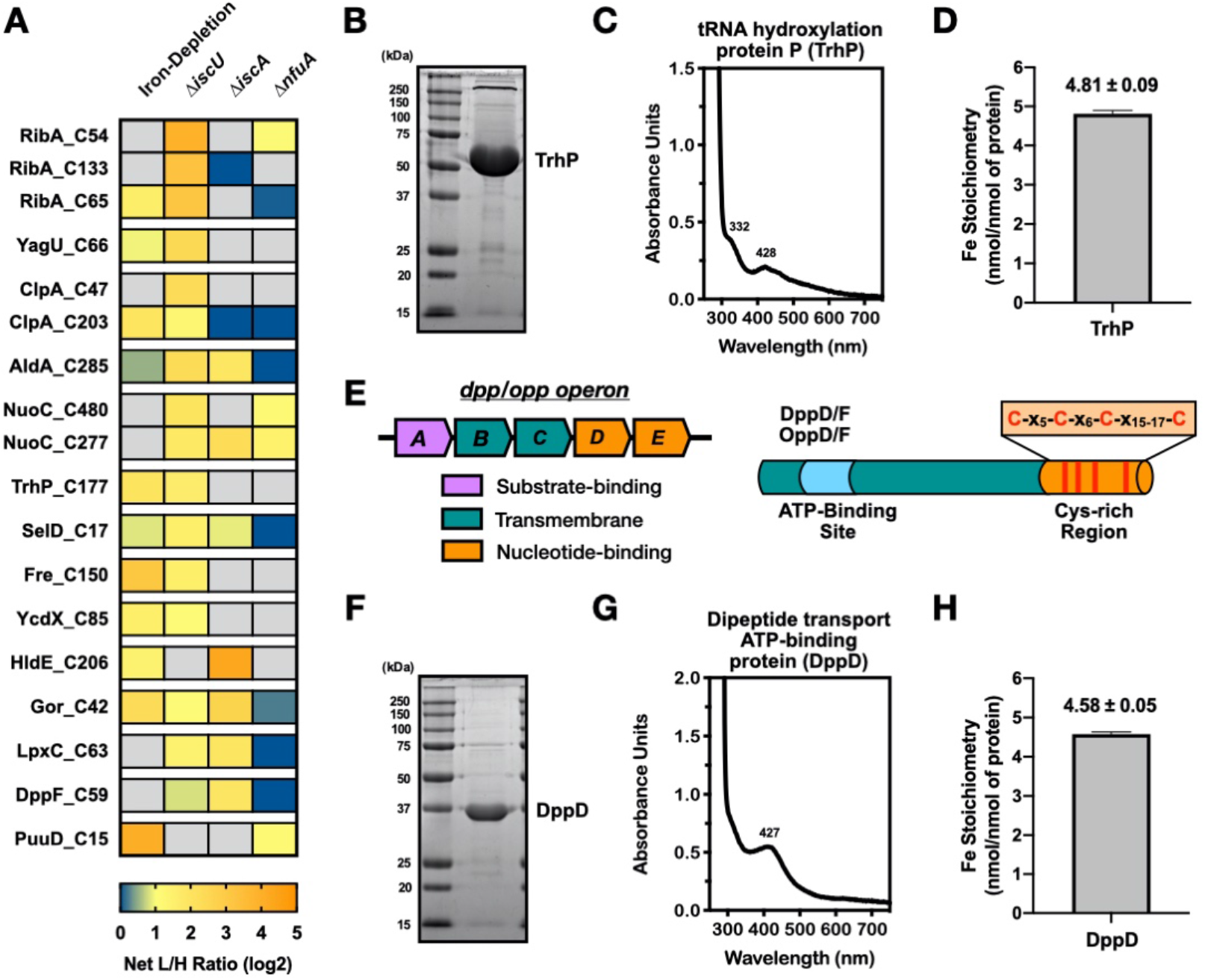
Predicting novel Fe-S binding sites from changes in cysteine reactivity. **a**, Heat map of putative Fe-S cluster ligands across all experimental growth conditions and cell strains. Only cysteine residues with net increases in cysteine reactivity > 1.5 in two experimental conditions were included (derived from the larger selection in **Supplemental Figure 6**). **b**, SDS-PAGE analysis of purified *E. coli* TrhP. **c**, UV-visible absorbance spectrum of purified TrhP. **d**, ICP-OES analysis of the iron content of purified TrhP. **e**, Organization of the *dpp* and *opp* operons and the protein domain structure of the nucleotide-binding subunits, DppD/F and OppD/F. Conserved cysteine residues in the C-terminal extension (orange) are shown as red lines. **f**, SDS-PAGE analysis of purified *E. coli* DppD. **g**, UV-visible absorbance spectrum of purified DppD. **h**, ICP-OES analysis of the iron content of purified DppD.

We then proceeded to determine if proteins on this list could be putative Fe-S cluster proteins by overexpression, purification, and subsequent analysis by UV-visible absorption spectroscopy and ICP-OES. The first protein that was assayed for Fe-S binding was TrhP (also called YegQ) (**Figure 5A**), which was recently identified as a tRNA hydroxylase involved in the generation of 5-hydroxyuridine.^47^ Recently, a close homolog of TrhP, RlhA, was shown to be similarly involved in ribosomal RNA hydroxylation. Interestingly, RNA hydroxylation levels decreased in an *isc* deletion strain, suggesting a dependence on the Fe-S biogenesis pathway.^48^ TrhP also has more distant homology to two Fe-S proteins involved in ubiquinone biosynthesis, UbiU and UbiV, but shows poor conservation of the putative cysteine ligands for the Fe-S cluster.^49^ The presence of a Fe-S cluster on TrhP or RlhA has yet to be confirmed. Here, we experimentally validated the presence of an Fe-S cluster on TrhP. TrhP was expressed, and purified from *E. coli* (**Figure 5B**), and the presence of an Fe-S cluster was observed by UV-visible absorption spectroscopy (**Figure 5C**) and ICP-OES (**Figure 5D**). TrhP likely binds a single [4Fe-4S] cluster due to the presence of ∼ 4-5 iron atoms per protein monomer by ICP-OES analysis.

A second putative Fe-S protein from our high-confidence list was DppF (**Figure 5A**). A close homolog of DppF, OppD, was also identified in our iron-depletion dataset and displayed a net increase in cysteine reactivity (**Supplemental Figure 6)**. These proteins are both nucleotide binding subunits of ABC transporter complexes.^50^ Both the DppABCDF and OppABCDF ABC transporters are involved in the energy dependent import of small peptides into cells. DppD/F and OppD/F contain a C-terminal extension that is rich in cysteine residues (**Figure 5E**). As a representative of this family of proteins, DppD was expressed and purified from *E. coli* (**Figure 5F**). The presence of an Fe-S cluster was confirmed by UV-visible absorption spectroscopy (**Figure 5G**), and ∼ 4-5 iron atoms per protein monomer was detected by ICP-OES (**Figure 5H**). Together, the experimental confirmation of the Fe-S cluster on TrhP and DppD serve to highlight the promise of our chemoproteomic platform to identify unknown Fe-S proteins by spotlighting cysteine residues with net increases in reactivity under conditions of dysregulated Fe-S biogenesis. Importantly, our approach to identify uncharacterized Fe-S proteins is completely unbiased and does not rely on sequence homology to previously identified Fe-S proteins, or Fe-S ligating cysteine motifs.

## Summary

A significant challenge in the field of Fe-S protein biology is the inability to monitor Fe-S cluster occupancy across multiple proteins within a native proteome. Typical methods to study Fe-S cluster binding include purification and spectroscopic analysis of individual proteins, or monitoring of radiolabeled iron incorporation by autoradiography. Here, we describe a chemoproteomic platform to systematically report on Fe-S occupancy within a proteome. This platform monitors the characteristic increase in the reactivity of Fe-S cluster cysteine ligands upon loss of the Fe-S cluster. Increases in cysteine reactivity are measured using the established isoTOP-ABPP platform, and coupled with protein abundance changes obtained by quantitative ReDiMe analyses, to provide net changes in cysteine reactivity across the proteome. Fe-S cysteine ligands show net increases in cysteine reactivity under conditions of dysregulated Fe-S biogenesis, including iron-depletion, and genetic deletion of *isc* Fe-S biogenesis components. This chemoproteomic platform provided deeper insight into Fe-S client proteins and the Fe-S biogenesis pathway including: (1) the demonstration that different types of Fe-S clusters (AdoMet radical versus auxiliary) within a single protein show differential occupancy under conditions of dysregulated Fe-S biogenesis (e.g. BioB, RimO); (2) the assignment of cysteine ligands for Fe-S cluster proteins where sites of cluster binding are uncharacterized (e.g. YdiJ); (3) the establishment that deletion of early components of Fe-S biogenesis (e.g. IscU) have significantly larger effects on the Fe-S proteome than deletion of late components (e.g. IscA and NfuA); (4) the delineation of client proteins dependent on the Isc and Suf Fe-S biogenesis pathways; (5) the differentiation of client Fe-S proteins between the IscA and NfuA A-type scaffold proteins; and, (6) the identification of previously unannotated Fe-S proteins in an unbiased manner (e.g. TrhP and DppD). In our analyses, we observed ∼70% coverage of Fe-S proteins and/or cysteine ligands from *E. coli*. Many of the unidentified members were ferredoxins that are selectively expressed under specialized growth conditions (e.g. anaerobic), and coverage will likely improve upon analysis of additional nutrient environments. Determining net changes in cysteine reactivity require the identification of a Fe-S protein in the unenriched proteomics (ReDiMe) analysis, as well as the quantification of cysteine labeling by isoTOP-ABPP. In many cases, we observed Fe-S proteins in one of these two analyses, likely due to the low abundance of some Fe-S proteins for ReDiMe analysis, or the lack of an ionizable IA-labeled peptide for isoTOP-ABPP analysis. The use of additional fractionation steps, as well as more sensitive mass-spectrometry instrumentation, will serve to alleviate some of these issues. Additionally, the identification of novel Fe-S proteins is complicated by the presence of secondary effects that could serve to increase cysteine reactivity, including conformational changes of a protein driven by loss of an interaction with an Fe-S protein binding partner. Therefore, significant biochemical and spectroscopic analysis needs to be performed to confirm the presence of an Fe-S cluster on putative Fe-S protein candidates. Other potential future applications of this platform include: (1) the characterization of client proteins served by other components of the Isc pathway (e.g. the accessory protein HscA); (2) the identification of unannotated Fe-S proteins under alternative growth conditions and in other bacterial strains; (3) the evaluation of the Fe-S biogenesis pathways in mammalian cells (mitochondrial and cytosolic Fe-S pathways); and, (4) the analysis of Fe-S cluster stability under cellular stress conditions (e.g. oxidative stress). We are confident that further improvement of our platform and expansion to other complex biological systems will provide a valuable tool to investigate Fe-S clusters within native proteomes, and continue to illuminate the biology of Fe-S clusters.

## Data Availability

Mass spectrometry data for all proteomic analyses will be made available upon request.

## Acknowledgements

We thank all members of the E.W. laboratory for discussions and feedback. This work was supported by NIH grant R35GM134964 to E.W.

## Author Contributions

D.W.B. and E.W. designed research; D.W.B. performed research and proteomic analysis; D.W.B. and E.W. analyzed and interpreted the data; D.W.B. and E.W. wrote the paper.

## Additional information

**Supplemental Information** for this paper is available in the online version

C**orrespondence** should be addressed to D.W.B. or E.W.

