## Supplemental Figures and Methods for "Monitoring iron-sulfur cluster occupancy across the *E. coli* proteome using chemoproteomics"

### Supplementary Figures

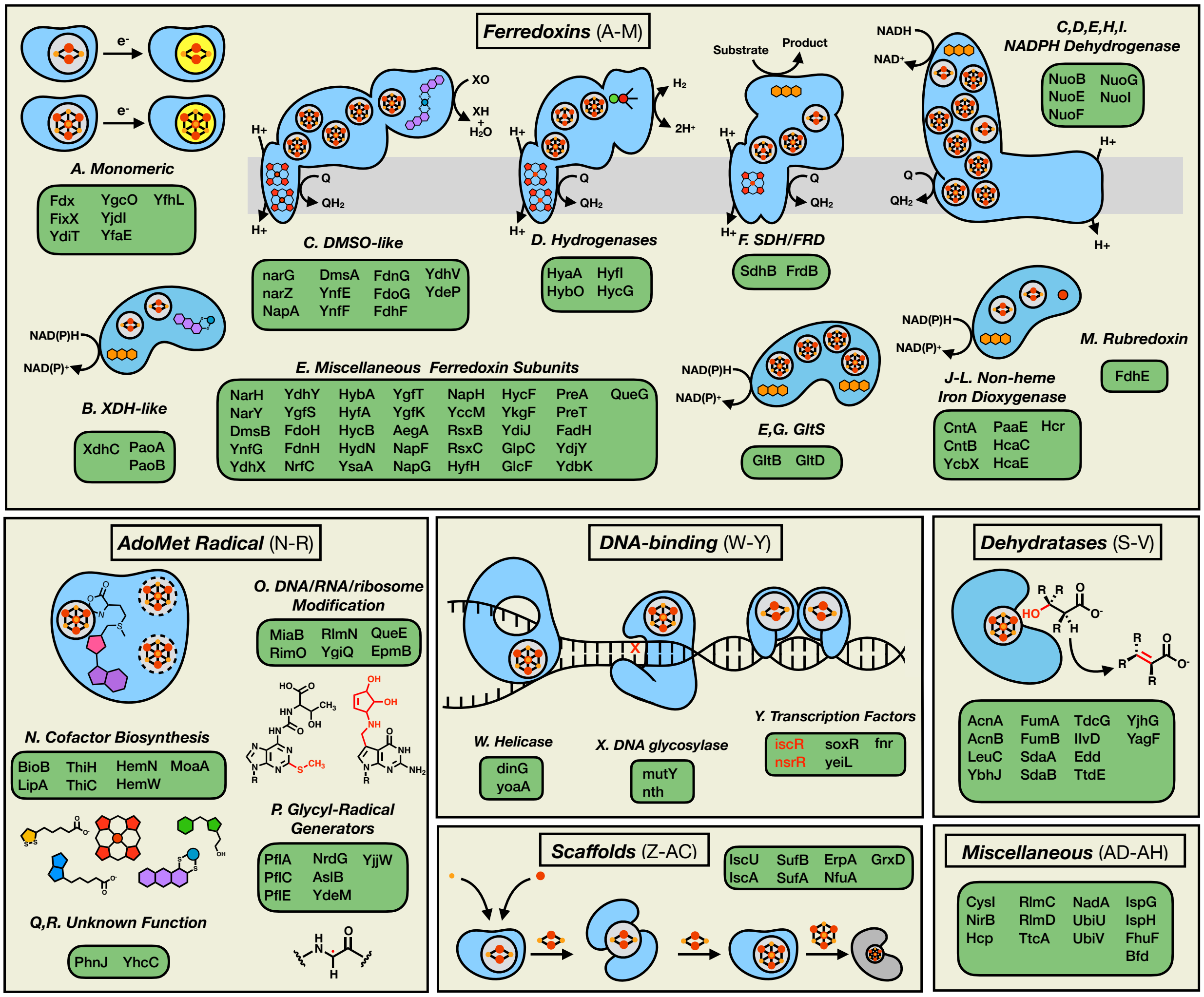

**Supplementary Figure 1 | A Functional catalogue of the *E. coli* Fe-S proteome.** The 144 members of the E. coli Fe-S proteome categorized by cluster function (see **Supplementary Table 1**)

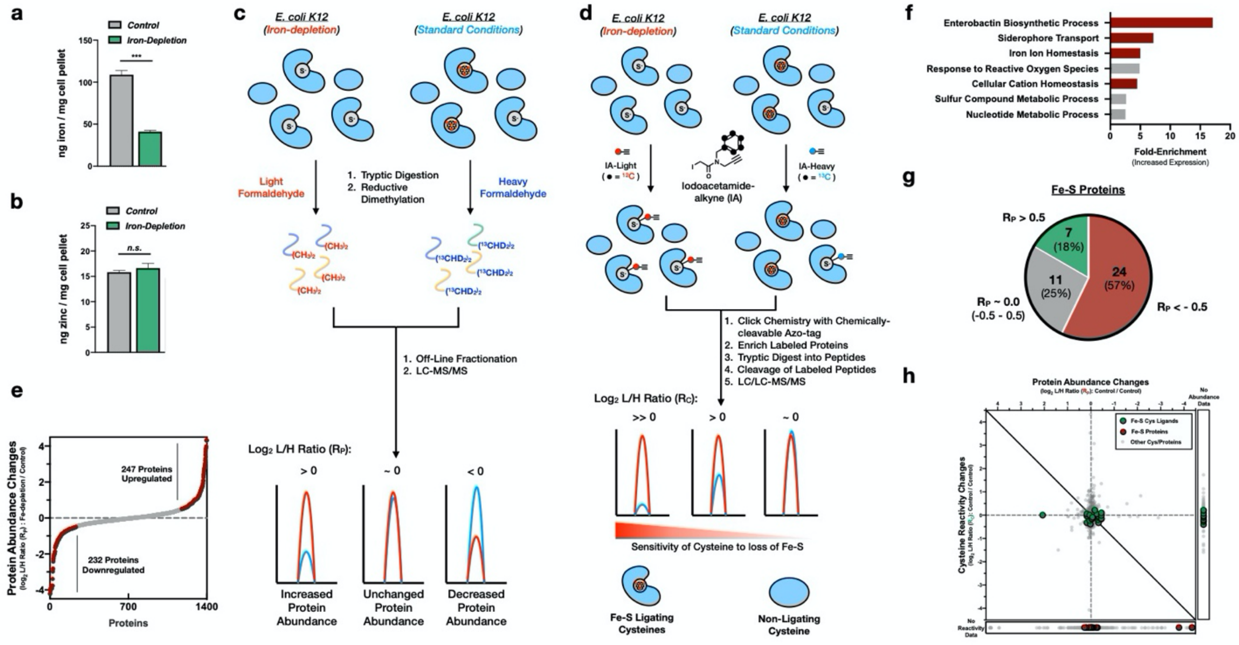

**Supplementary Figure 2 | Cysteine labeling and protein abundance changes in the *E. coli* Fe-S proteome upon iron-depletion. a,b,** Intracellular iron (**a**) and zinc (**b**) concentration determined by ICP-OES for *E. coli* grown under iron-replete (control) (gray) or iron-depleted (green) conditions. Significance is calculated as *** $p < 0.005$, paired t-test (two-tailed) with $n = 3$. Error bars represent the standard error of the mean. **c,** Reductive dimethylation (ReDiMe) proteomic platform employed to measure protein abundance changes upon growth of *E. coli* under iron-depletion conditions. **d,** IsoTOP-ABPP proteomic platform employed to measure cysteine labeling changes upon growth of *E. coli* under iron-depletion conditions. **e,** Waterfall plot of $\log_2$ L/H ratio ($R_P$) changes for all quantified proteins from an iron-depleted *E. coli* proteome (see **Supplemental Table 2**). Proteins with $R_P > |0.5|$ are highlighted in red. **f,** Gene-ontology analysis of processes enriched in proteins with $R_P > 0.5$. Processes involved in iron import and homeostasis are highlighted in red. **g,** Pie chart analysis of the number of quantified Fe-S proteins with $R_P < -0.5$ (red), $R_P \sim 0.0$ (gray), and $R_P > 0.5$ (green). **h,** Two-dimensional proteomic dataset for the *E. coli* proteome grown under iron-replete (control) conditions (see **Supplemental Table 3**). All quantified cysteine residues are plotted in the main graph (annotated Fe-S cluster cysteine ligands - green circles, non-annotated cysteine residues - light gray small circles). Inset to right: cysteine residues with no protein abundance data (annotated Fe-S cluster cysteine ligands - green circles, non-annotated cysteine residues - light gray small circles). Inset below: proteins with no cysteine reactivity data (annotated Fe-S protein - red circles, non-annotated proteins - light gray small).

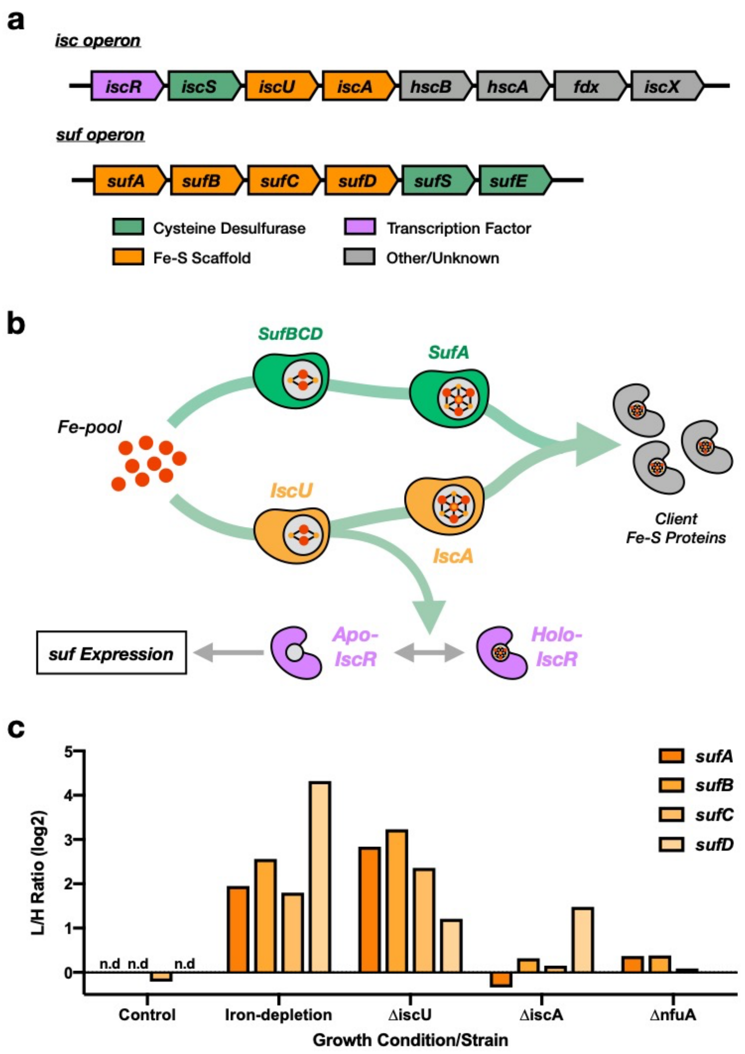

**Supplementary Figure 3 | *suf* operon expression in *E. coli isc* genetic deletion strains. a,** Structural organization of the *E. coli isc* and *suf* operons. **b**, Proposed model of *suf* operon function and regulation by the IscU client protein and transcription factor, IscR. **c**, Protein abundance changes for scaffold complex proteins from the *suf* operon across all experimental growth conditions and deletion strains.

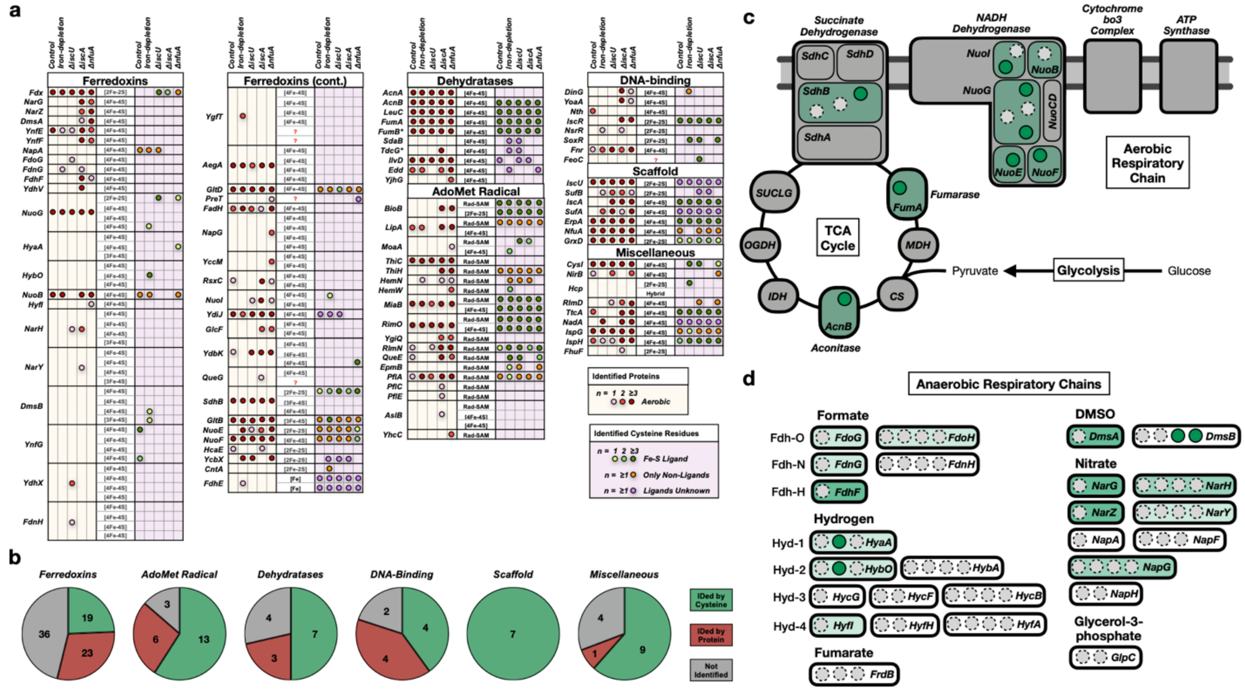

**Supplementary Figure 4 | Map of cysteine and protein coverage across the *E. coli* Fe-S proteome. a,** Table of identified Fe-S proteins (orange cells) and clusters (purple cells) across all growth conditions and deletion strains. Identification is indicated by a closed circle, for either protein abundance (red circles) or cysteine reactivity (annotated Fe-S cluster cysteine ligands - green circles, non-Fe-S ligating cysteine residues – orange circles, cysteine residues, where Fe-S ligands are unknown for the protein – purple circle). **b,** Pie chart indicating the proteomic coverage for each functional class of Fe-S proteins. Proteins are grouped by identification through cysteine reactivity (green), protein abundance (red), or proteins not identified in this study (gray). **c,** Diagram of Fe-S protein and cluster identification during aerobic respiration on glucose as a carbon source. Identified Fe-S proteins are shaded in green, while identified cluster are indicated in closed green circles. Clusters that were not identified are indicated by dashed gray circles and non-Fe-S proteins are colored gray. **d**, Diagram of Fe-S protein and cluster identification in anaerobic respiration complexes with alternative electron acceptors and donors. Identified Fe-S proteins are shaded in green, while identified cluster are indicated in closed green circles. Clusters that were not identified are indicated by dashed gray circles.

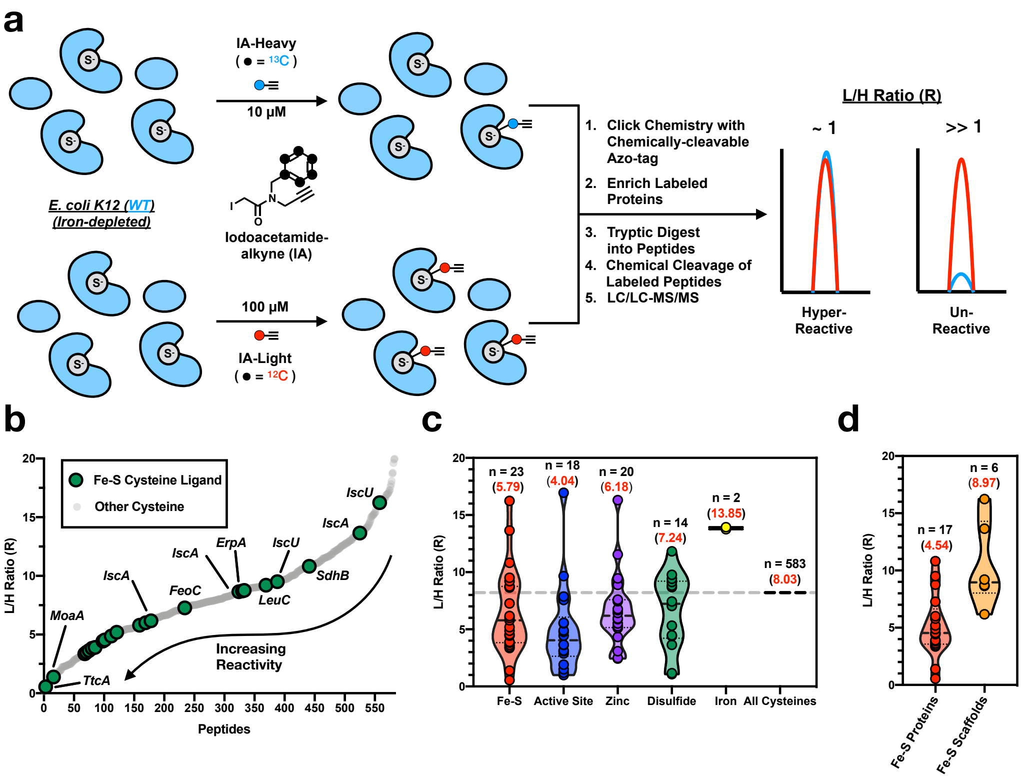

**Supplementary Figure 5 | Reactivity profile of cysteine residues involved in Fe-S ligation. a,** Proteomic workflow for measuring the cysteine reactivity of Fe-S cluster ligands in an iron-depleted *E. coli* proteome labeled with 100 μM IA-Light or 10 μM IA-Heavy. **b,** Cysteine reactivity curve for all quantified cysteine residues (light gray) from an iron-depleted *E. coli* proteome (see **Supplemental Table 7**). Fe-S cluster cysteine ligands are highlighted (green circles). **c,d** Violin plot of L/H ratios (R) for unique groups of functional cysteine residues (**c**), including Fe-S ligands (red), active site residues (blue), zinc ligands (purple), disulfides (green), and iron ligands (yellow) and for Fe-S ligands from Fe-S client proteins (red) and Fe-S scaffold proteins (yellow) (**d**). The median R value for each functional group of cysteine residues is displayed as a dashed line, while the average R values (red) and number of unique values (black) are indicated for each group.

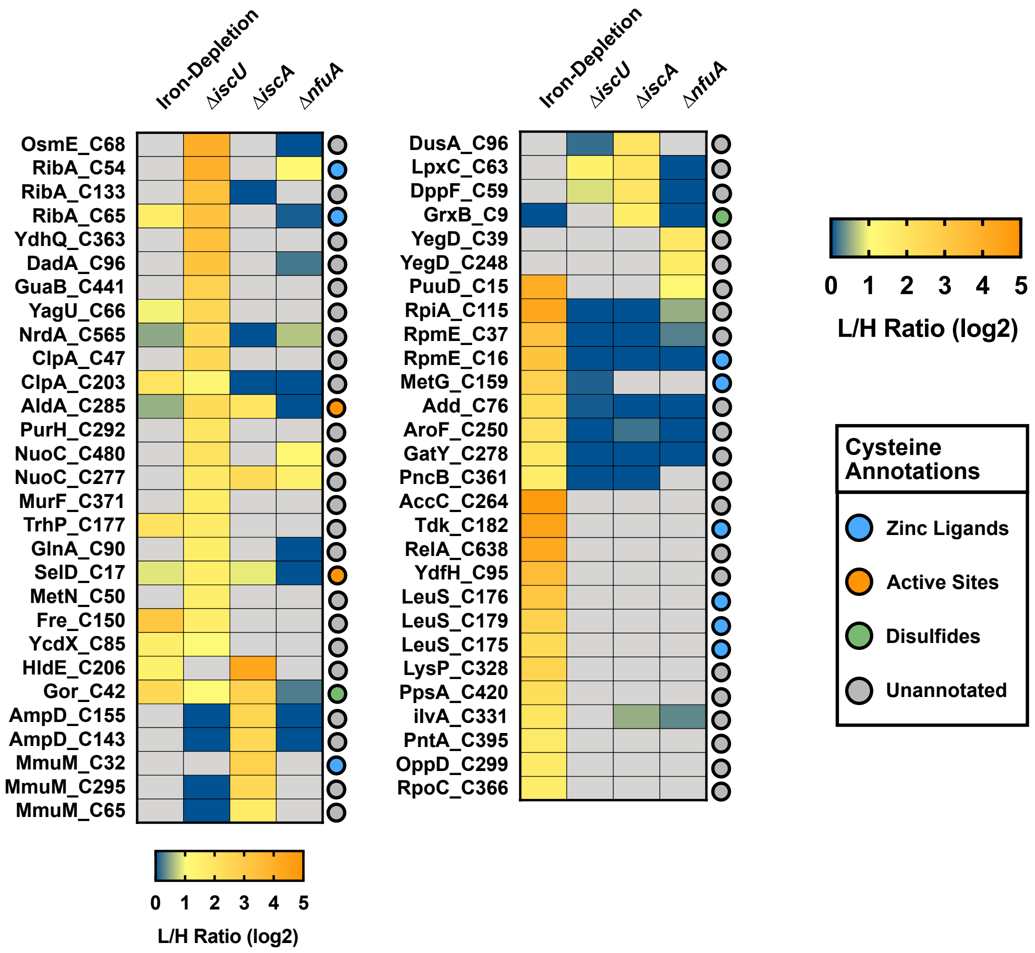

**Supplementary Figure 6 | Putative Fe-S binding sites based on net changes in cysteine reactivity upon perturbation of Fe-S biogenesis.** Heat map of putative Fe-S cluster cysteine ligands across all experimental growth conditions and deletion strains with net changes in cysteine reactivity ($log_2$ L/H ratios > 1.5) in at least one dataset. Known functional annotations for each cysteine residue are indicated to the right; zinc ligand – blue, active site – orange, disulfide – green, unannotated – gray.

### Supplementary Tables

**Supplementary Table 1 | The *E. coli* Fe-S proteome.** Catalogue of Uniprot annotated *E. coli* Fe-S proteins, separated into functional and sequence similarity related groups. The Fe-S cluster type and the protein-based ligands are indicated, when they are known.

**Supplementary Table 2 | Quantitative two-dimensional chemoproteomic dataset for *E. coli* grown under iron-depletion conditions. a-b,** raw ReDiMe L/H (**a**) and H/L (**b**) peptide ratios calculated using cimage software and grouped by protein for WT *E. coli* K12 cells grown under iron-depleted (light) or iron-replete (heavy) conditions (n = 4 biologically independent samples). **c,** full dataset of quantitative protein abundance changes reported as average $\log_2$ L/H ratios (L/H and H/L ratios for each protein collated, corrected to the median, and averaged from **a** and **b**). **d-e,** raw IA-alkyne modified L/H (**d**) and H/L (**e**) cysteine-containing peptide ratios calculated using cimage software for WT *E. coli* K12 cells grown under iron-depleted (light) or iron-replete (heavy) conditions (n = 4 biologically independent samples). **f,** full dataset of quantitative cysteine reactivity changes reported as average $\log_2$ L/H ratios (L/H and H/L ratios for each peptide collated, corrected to the median, and averaged from **d** and **e**). **g**, Corrected $\log_2$ L/H ratios, for cysteine containing peptides (from **f**) that also had reported protein abundance ratios (from **c**). **h**, $\log_2$ L/H ratios, for cysteine containing peptides (from **f**) that did not have reported protein abundance ratios. **i**, $\log_2$ L/H ratios, for proteins (from **c**) that did not have any reported cysteine peptide ratios.

**Supplementary Table 3 | Quantitative two-dimensional chemoproteomic dataset for *E. coli* grown under iron-replete (control) conditions. a-b,** raw ReDiMe L/H (**a**) and H/L (**b**) peptide ratios calculated using cimage software and grouped by protein for WT *E. coli* K12 cells grown under iron-replete (light) or iron-replete (heavy) conditions (n = 3 biologically independent samples). **c,** full dataset of quantitative protein abundance changes reported as average $\log_2$ L/H ratios (L/H and H/L ratios for each protein collated, corrected to the median, and averaged from **a** and **b**). **d-e,** raw IA-alkyne modified L/H (**d**) and H/L (**e**) cysteine-containing peptide ratios calculated using cimage software for WT *E. coli* K12 cells grown under iron-replete (light) or iron-replete (heavy) conditions (n = 3 biologically independent samples). **f,** full dataset of quantitative cysteine reactivity changes reported as average $\log_2$ L/H ratios (L/H and H/L ratios for each peptide collated, corrected to the median, and averaged from **d** and **e**). **g**, Corrected $\log_2$ L/H ratios, for cysteine containing peptides (from **f**) that also had reported protein abundance ratios (from **c**). **h**, $\log_2$ L/H ratios, for cysteine containing peptides (from **f**) that did not have reported protein abundance ratios. **i**, $\log_2$ L/H ratios, for proteins (from **c**) that did not have any reported cysteine peptide ratios.

**Supplementary Table 4 | Quantitative two-dimensional chemoproteomic dataset for *E. coli* ∆*iscU* grown under iron-replete conditions. a-b,** raw ReDiMe L/H (**a**) and H/L (**b**) peptide ratios calculated using cimage software and grouped by protein for *E. coli* K12 ∆*iscU* cells grown under iron-replete (light) or for *E. coli* K12 WT cells grown under iron-replete (heavy) conditions (n = 3 biologically independent samples). **c,** full dataset of quantitative protein abundance changes reported as average $log_2$ L/H ratios (L/H and H/L ratios for each protein collated, corrected to the median, and averaged from **a** and **b**). **d-e,** raw IA-alkyne modified L/H (**d**) and H/L (**e**) cysteine-containing peptide ratios calculated using cimage software for *E. coli* K12 Δ*iscU* cells grown under iron-replete (light) or for *E. coli* K12 WT cells grown under iron-replete (heavy) conditions (n = 5 biologically independent samples). **f,** full dataset of quantitative cysteine reactivity changes reported as average log2 L/H ratios (L/H and H/L ratios for each peptide collated, corrected to the median, and averaged from **d** and **e**). **g**, Corrected $log_2$ L/H ratios, for cysteine containing peptides (from **f**) that also had reported protein abundance ratios (from **c**). **h**, $log_2$ L/H ratios, for cysteine containing peptides (from **f**) that did not have reported protein abundance ratios. **i**, $log_2$ L/H ratios, for proteins (from **c**) that did not have any reported cysteine peptide ratios.

**Supplementary Table 5 | Quantitative two-dimensional chemoproteomic dataset for *E. coli* Δ*iscA* grown under iron-replete conditions. a-b,** raw ReDiMe L/H (**a**) and H/L (**b**) peptide ratios calculated using cimage software and grouped by protein for *E. coli* K12 Δ*iscA* cells grown under iron-replete (light) or for *E. coli* K12 WT cells grown under iron-replete (heavy) conditions (n = 4 biologically independent samples). **c,** full dataset of quantitative protein abundance changes reported as average $log_2$ L/H ratios (L/H and H/L ratios for each protein collated, corrected to the median, and averaged from **a** and **b**). **d-e,** raw IA-alkyne modified L/H (**d**) and H/L (**e**) cysteine-containing peptide ratios calculated using cimage software for *E. coli* K12 Δ*iscA* cells grown under iron-replete (light) or for *E. coli* K12 WT cells grown under iron-replete (heavy) conditions (n = 4 biologically independent samples). **f,** full dataset of quantitative cysteine reactivity changes reported as average $log_2$ L/H ratios (L/H and H/L ratios for each peptide collated, corrected to the median, and averaged from **d** and **e**). **g**, Corrected $log_2$ L/H ratios, for cysteine containing peptides (from **f**) that also had reported protein abundance ratios (from **c**). **h**, $log_2$ L/H ratios, for cysteine containing peptides (from **f**) that did not have reported protein abundance ratios. **i**, $log_2$ L/H ratios, for proteins (from **c**) that did not have any reported cysteine peptide ratios.

**Supplementary Table 6 | Quantitative two-dimensional chemoproteomic dataset for *E. coli* Δ*nfuA* grown under iron-replete conditions. a-b,** raw ReDiMe L/H (**a**) and H/L (**b**) peptide ratios calculated using cimage software and grouped by protein for *E. coli* K12 Δ*nfuA* cells grown under iron-replete (light) or for *E. coli* K12 WT cells grown under iron-replete (heavy) conditions (n = 4 biologically independent samples). **c,** full dataset of quantitative protein abundance changes reported as average $log_2$ L/H ratios (L/H and H/L ratios for each protein collated, corrected to the median, and averaged from **a** and **b**). **d-e,** raw IA-alkyne modified L/H (**d**) and H/L (**e**) cysteine-containing peptide ratios calculated using cimage software for *E. coli* K12 Δ*nfuA* cells grown under iron-replete (light) or for *E. coli* K12 WT cells grown under iron-replete (heavy) conditions (n = 4 biologically independent samples). **f,** full dataset of quantitative cysteine reactivity changes reported as average $log_2$ L/H ratios (L/H and H/L ratios for each peptide collated, corrected to the median, and averaged from **d** and **e**). **g**, Corrected $log_2$ L/H ratios, for cysteine containing peptides (from **f**) that also had reported protein abundance ratios (from **c**). **h**, $log_2$ L/H ratios, for cysteine containing peptides (from **f**) that did not have reported protein abundance ratios. **i**, $log_2$ L/H ratios, for proteins (from **c**) that did not have any reported cysteine peptide ratios.

**Supplementary Table 7 | Cysteine reactivity dataset for the *E. coli* proteome. a,** raw IA-alkyne modified L/H cysteine-containing peptide ratios calculated using cimage software for WT *E. coli* K12 cell lysates labeled with either 100 μM IA-alkyne (light) or 10 μM IA-alkyne (heavy). **b**, full cysteine reactivity dataset (from **a**) of average peptide L/H ratios (n = 3 biologically independent samples). **c**, filtered cysteine reactivity data set (from **b**), with all singleton (R = 20) values removed.

**Supplementary Table 8 | E. coli strains used in this study.**

| Strain | Organism | ID | Genotype |
|---|---|---|---|
| Control | *E. coli* | BW25113 | F-, *Δ(araD-araB)567*, *ΔlacZ4787*(::rrnB-3), *λ*$^-$, *rph-1*, *Δ(rhaD-rhaB)568*, *hsdR514* |
| *ΔiscU* | *E. coli* | JW2513-1 | F-, *Δ(araD-araB)567*, *ΔlacZ4787*(::rrnB-3), *λ*$^-$, *ΔiscU775::kan*, *rph-1*, *Δ(rhaD-rhaB)568*, *hsdR514* |
| *ΔiscA* | *E. coli* | JW2512-1 | F-, *Δ(araD-araB)567*, *ΔlacZ4787*(::rrnB-3), *λ*$^-$, *ΔiscA774::kan*, *rph-1*, *Δ(rhaD-rhaB)568*, *hsdR514* |
| *ΔnfuA* | *E. coli* | JW3377-7 | F-, *Δ(araD-araB)567*, *ΔlacZ4787*(::rrnB-3), *λ*$^-$, *ΔgntY748::kan*, *rph-1*, *Δ(rhaD-rhaB)568*, *hsdR514* |

**Supplementary Table 9 | Primers used in this study.**

| Gene name | Uniprot ID | Strain | Forward Primer | Reverse primer |
|---|---|---|---|---|
| *acnA* | P25516 | *E. coli (*K12) | TATCGAAGGTCGTCATATGTCGTCAACCCTACGAGAAG | TTAGCAGCCGGATCCTCGAGTTATTCAACATATTACGAATGACATAATGC |
| *trhP* | P76403 | *E. coli (*K12) | TATCGAAGGTCGTCATATGTTTAAACCGGAACTCCTTTCC | TTAGCAGCCGGATCCTCGAGTCACTTACCGTGGGGATTA |
| *dppD* | P0AAG0 | *E. coli (*K12) | TATCGAAGGTCGTCATATGGCGTTATTAAATGTAGATAAATTATCGGT | TTAGCAGCCGGATCCTCGAGTCATAGTGTCGGCCTCCC |

### Experimental Procedures

***E. coli* strains and growth media.** *E. coli* mutant strains were obtained from the Keio collection (**Supplemental Table 8)**.[1] These strains are derived from the *E. coli* K-12 strain BW25113.[2] All growths were carried out in MOPS-minimal media; standard (iron-replete) recipe - 40 mM MOPS and 4 mM tricine (adjusted to pH 7.4 with KOH), 0.1 M NaCl, 10 mM $NH_4Cl$, 1.32 mM $KH_2PO_4$, 0.523 mM $MgCl_2$, 0.276 mM $Na_2SO_4$, 0.1 mM $FeSO_4$, trace micronutrients, and 0.1% glucose.[3] Iron-depleted MOPS-minimal media was prepared as the standard recipe with the exclusion of supplemental iron. The *E. coli* BL21(DE3) strain was used for all protein over-expression experiments. All Glassware used for culturing *E. coli* was acid washed to remove any residual metals.

***E. coli* Fe-S protein cloning and expression.** The full-length gene sequences for the *E. coli* Fe-S proteins AcnA, TrhP, and DppD were PCR amplified from *E. coli* K-12 genomic DNA with the appropriate primers (**Supplemental Table 9**). These sequences were cloned by Gibson assembly[4] into the pET-16b vector (double digested with the restriction enzymes *XhoI* and *NdeI*). These constructs generate a recombinant target protein with an N-terminal 6x His-tag. The resulting plasmids were verified by DNA sequencing and transformed into BL21(DE3) cells. For AcnA, transformed cells were cultured (37 ºC) in either iron-replete and iron-depleted MOPS-minimal media, supplemented with ampicillin (50 mg $l^{-1}$). Once an optical density at 600 nm ($OD_{600}$) of ~0.6 was reached, protein production was induced (37 ºC, 3 hr) by the addition of 0.4 mM IPTG. For TrhP and DppD, transformed cells were culture (37 ºC) in 2xYT media, supplemented with $FeSO_4^{-2}$ (1 g $l^{-1}$) and ampicillin (50 mg $l^{-1}$). Once an $OD_{600}$ of ~0.6 was reached, protein production was induced (25 ºC, 16hr) by the addition of 0.1 mM IPTG. For all three Fe-S proteins, cells were pelleted from 500 mL cultures by centrifugation (5,000 *g*, 10 min, 4 ºC).

**Purification and characterization of recombinant AcnA**. Cell pellets were resuspended in 5 mL lysis buffer (5 mM imidazole in PBS), and lysed by sonication (3 x10 pulses – 90%) on ice. Cell lysates were centrifuged (10,000 *g*, 10 min, 4 ºC) to remove unlysed cell debris. The resulting supernatant was loaded onto 1 mL of Ni-NTA that had been equilibrated with lysis buffer. The Ni-NTA was washed with 5-bed volumes wash buffer (25 mM imidazole in PBS). Recombinant AcnA was eluted from the Ni-NTA column with 2-bed volumes of elution buffer (250 mM imidazole in PBS). The protein was desalted over a pre-packed PD-10 column equilibrated with PBS buffer and concentrated using an Amicon-Ultra 10 kDa centrifugal filter. Protein concentration was determined by Bradford assay, and the presence or absence of an Fe-S cluster was confirmed by UV-visible absorption spectroscopy on a ThermoScientific Nanodrop 2000c spectrophotometer.

**IA-alkyne labeling for fluorescent gel analysis of AcnA.** Fifty micrograms of Recombinant AcnA (1 mg/ml), purified from growth in iron-replete or iron-depleted MOPS-minimal media, was incubated with 100 µM IA-alkyne (10 mM stock in DMSO) for 1 hr at 25 ºC. Click chemistry was performed with 25 µM rhodamine-azide (1.25 mM stock in DMSO), 1 mM TCEP, 100 µM TBTA ligand (17x stock in DMSO:t-butanol 1:4) and 1 mM $CuSO_4$ (50x stock in water). The reaction was quenched after 1 hr at 25 ºC by the addition of 50 µL of 2x SDS-PAGE loading buffer. Samples were separated by SDS-PAGE, visualized for IA-alkyne labeling by fluorescence on a BioRad ChemiDoc MP Gel Imager, stained with Coomassie-blue, and imaged again for protein abundance.

***E. coli* cell pellet preparation for ICP-OES.** The control *E. coli* K-12 strain, BW25113, was streaked out on an LB agar plate from a glycerol stock and incubated overnight at 37 ºC. The following day, single colonies were selected to inoculate 5 mL of iron-replete MOPS-minimal media, which were then incubated overnight at 37 ºC. Overnight cultures were used to inoculate 100 mL cultures of either iron-replete or iron-deplete MOPS-minimal media at a 1:100 dilution and grown at 37 ºC to an $OD_{600}$ of ~0.6. Cells were pelleted by centrifugation (5,000 *g*, 10 min, 4 ºC), washed twice with 500 µL of 1 mM EDTA in PBS and twice more with 500 µL of EDTA-free PBS. Cell pellets were dried down overnight on a speed-vac, before the dry pellet weight was determined. Dried cell pellets were dissolved overnight in 150 µL of TraceMetal Grade Nitric Acid, then heated for 1 hr at 80 ºC. The sample was then diluted to 5% nitric acid with Milli-Q double distilled water. Iron and zinc content was measured on an Agilent 5100 ICP-OES. Samples were prepared in triplicate and the relative iron and zinc concentration of each cell pellet sample was calculated from a calibration curve of external standards (iron – 0 to 56 mg $l^{-1}$, zinc – 0 to 65 mg $ml^{-1}$) according to peak height.

***E. coli* lysate preparation for two-dimensional chemoproteomics.** *E. coli* K-12 strains were streaked out onto LB agar plates from glycerol stocks and incubated overnight at 37 ºC. The following day, single colonies were selected to inoculate 5 mL of iron-replete MOPS-minimal media, which were then incubated overnight at 37 ºC. Overnight cultures were used to inoculate 100 mL cultures of either iron-replete or iron-deplete MOPS-minimal media at a 1:100 dilution and grown at 37 ºC to an $OD_{600}$ of ~0.6. Cells were pelleted by centrifugation (5,000 *g*, 10 min, 4 ºC) and washed once with 5 mL of PBS. Cell pellets were then resuspended in 500 µL of PBS, lysed by sonication and centrifuged (10,000 *g*, 10 min, 4 ºC) to generate clarified lysates. Protein concentration was determined by Bradford assay and lysates were diluted to a concentration of 2 mg/mL with PBS.

**Whole-proteome sample preparation for ReDiMe analysis.** For quantification of protein abundance changes upon iron-depletion or in strains harboring *isc* deletions (**Supplemental Tables 2-6**), 100 µg of both a control (iron-replete *E. coli* (K-12)) and Fe-S deficient (iron-depleted *E. coli* (K-12), *ΔiscU*, *ΔiscA*, or Δ*nfuA*) proteome (100 µL, 1 mg/mL) were precipitated by the addition of 5 µL 100% trichloroacetic acid in PBS, vortexed and frozen (-80 ºC, overnight). After thawing, proteins were pelleted by centrifugation (15,000 g, 10 min, 4 ºC) and the solvent was removed. Protein pellets were resuspended in 500 µL of ice-cold acetone by bath sonication and pelleted by centrifugation (5000 g, 10 min, 4 ºC). The solvent was removed and the pellet was allowed to air dry before being resuspended in 8 M Urea in 100 mM TEAB (30 µL). Reductive alkylation was performed by the sequential addition of 100 mM TEAB (70 µL), 1.5 µL of 1 M DTT (65 ºC, 15 min), and 2.5 µL of 400 mM iodoacetamide (25 ºC, 30 min). Reactions were diluted with additional 100 mM TEAB (120 µL) and tryptic digestion performed by the addition of 2 µg of sequencing-grade trypsin (4 µL of 20 µg diluted in $H_2O$) and 2.5 µL of 100 mM $CaCl_2$ (37 ºC, overnight). After tryptic digest, reductive dimethylation[5] was performed by the addition of 4 µL of 20% light (Fe-S deficient) or heavy (control) formaldehyde and 20 µL of 0.6 M sodium cyanoborohydride (25 ºC, 2 hours). The reaction was quenched by the addition of 8µL ammonium hydroxide (25 ºC, 15 min). The light (Fe-S deficient) and heavy (control) tryptic peptide samples were then combined, desalted on a Sep-Pak, and dried by speed-vac. Peptide samples were stored at -20 ºC until ready for fractionation and LC-MS/MS analysis.

**Off-line high-pH ReDiMe peptide fractionation.** Samples were resuspended in 500 µL of high pH buffer A (95% $H_2O$, 5% acetonitrile, 10 mM ammonium bicarbonate) and loaded onto a manual injection loop connected to an Agilent 1100 Series HPLC. Peptides were separated on a 25 cm Agilent Extend-C18 column using a 60 min gradient from 20-35% high pH buffer B (10% H2O, 90% acetonitrile, 10 mM ammonium bicarbonate).[6] Fractions were collected using a Gilson FC203B fraction collector into a 96 deep-well plate (0.6 min/well). Subsequent concatenation of every sixth well resulted in six pooled fractions for later LC-MS/MS analysis. These six fractions were dried by speed-vac and then resuspended in 30 µL of low pH buffer A (95% $H_2O$, 5% acetonitrile, 0.1% formic acid).

**LC-MS/MS and data analysis for ReDiMe whole-proteome quantification.** LC-MS/MS was performed on a Thermo LTQ Oribtrap XL mass spectrometer (Thermo Scientific) coupled to an easy-nanoLC system (Agilent). For each of the six off-line fractions, 10 µL of peptide mixture was pre-loaded onto a C18 pre-column. Peptides were eluted onto a 100-mm fused silica column packed with 10 cm C18 resin using a 4 hr gradient of buffer B in Buffer A (Buffer A: 95% water, 5% acetonitrile, 0.1% formic acid; Buffer B: 20% water, 80% acetonitrile, 0.1% formic acid). The flow rate through the column was ~0.4 µL/min, with a spray voltage of 2.75 kV. One full MS1 scan (400-1800 MW) was followed by 8 data dependent scans of the $n^{th}$ most intense ion. Dynamic exclusion was enabled. The tandem MS data was analyzed by the SEQUEST algorithm.[7] Static modification of cysteine residues (+57.0215 m/z, iodoacetamide alkylation) was assumed with no enzyme specificity. The precursor-ion mass tolerance was set at 50 ppm while the fragment-ion mass tolerance was set to 0 (default setting). Data was searched against an *E. coli* reverse-concatenated non-redundant FASTA database containing Uniprot identifiers. MS datasets were independently searched with light and heavy ReDiMe parameter files; for these searches, static modifications on lysine and the peptide N-terminus of +28.0313 (light) or +34.0632 (heavy) were used. MS2 spectra matches were assembled into protein identifications and filtered using DTASelect2.0,[8] to generate a list of protein hits with a peptide false-discovery rate of <5%. Quantification of protein L/H ratios were calculated using the cimage quantification package described previously.[9]

**Chemoproteomic labeling and enrichment of E. coli cysteine residues.** For identification and enrichment of cysteines residues labeled upon iron-depletion or in strains harboring *isc* deletions (**Supplemental Tables 2-6**), 2 mg of control (iron-replete *E. coli* (K-12)) or Fe-S deficient (iron-depleted *E. coli* (K-12), *ΔiscU*, *ΔiscA*, or Δ*nfuA*) proteome (1 mL, 2 mg/mL) were labeled with 100 µM IA-alkyne light (IAL – 10 µL, 10 mM stock in DMSO) or IA-alkyne Heavy (IAH – 10 µL, 10 mM stock in DMSO) for 1 h at room temperature, respectively. Probe labeled proteins were then conjugated to chemically-cleavable diazobenzene biotin-azide (azo-tag) by Cu(I)-catalyzed [3 + 2] cycloaddition as previously reported.[10,11] Briefly, azo-tag (100 µM), TCEP (1 mM), TBTA (100 µM), and $CuSO_4$ (1 mM) were sequentially added to the labeled proteome. Reactions were vortexed and incubated at room temperature for 1 h. Light- and heavy-proteomes were combined and centrifuged (6500 *g*, 10 min, 4 °C) to collect precipitated protein. Supernatant was discarded, and protein pellets were resuspended in 500 µL of methanol (dry-ice chilled) with sonication, and centrifuged (6500 *g*, 10 min, 4 °C). This step was repeated, and the resulting washed pellets was redissolved (1.2% w/v SDS in PBS; 1 mL); sonication followed by heating at 80-95 °C for 5 min was used to ensure complete solubilization. Samples were cooled to room temperature, diluted with PBS (5.0 mL), and incubated with Streptavidin beads (100 µL of 50% aqueous slurry per enrichment) overnight at 4 °C. Samples were allowed to warm to room temperature, pelleted by centrifugation (1400 *g*, 3 min), and supernatant discarded. Beads were then sequentially washed with 0.2% SDS in PBS (5 mL x 1), PBS (5 mL x 3) and H2O (5 mL x 3) for a total of 7 washes.

**On-bead reductive alkylation, tryptic digestion, and chemical cleavage of samples.** Following the final wash, protein-bound streptavidin beads were resuspended in 6 M urea in PBS (500 µL) and reductively alkylated by sequential addition of 10 mM DTT (25 µL of 200 mM in H2O, 65 °C for 20 min) and 20 mM iodoacetamide (25 µL of 400 mM in H2O, 37 °C for 30 min) to each sample. Reactions were then diluted by addition of PBS (950 µL), pelleted by centrifugation (1400 *g*, 3 min), and the supernatant discarded. Samples were then subjected to tryptic digestion by addition of 200 µL of a pre-mixed solution of 2M urea in PBS, 1 mM CaCl2 (2 µL of 100 mM in H2O), and 2 µg of Trypsin Gold (Promega, 4 µL of 0.5 µg/µL in 1% acetic acid). Samples were shaken overnight at 37 °C and pelleted by centrifugation (1400 *g*, 3 min). Beads were then washed sequentially with PBS (500 µL x 3) and H2O (500 µL x 3), and then resuspended in sodium dithionite (50 µL of 50 mM in $H_2O$) and incubate at room temperature for 1 h to release labeled peptides. Beads were pelleted by centrifugation (1,400 g, 3 min, 25 $^oC$) and the supernatant transferred to a new centrifuge tube. Cleavage was repeated twice more (75 µL of 50 mM in $H_2O$) with the supernatants transferred to same centrifuge tube as the previous cleavage supernatant. The beads were washed with PBS (100 µL, 2x), with the washes being combined with the previous supernatants, for a total eluted peptide sample volume of ~ 350 µL. Formic acid (17.5 µL) was added to a final concentration of 5% and samples were stored at -20 $^oC$ until ready for LC/LC-MS/MS analysis.

**LC/LC-MS/MS and data analysis for isoTOP-ABPP.** Mass spectrometry was performed using a Thermo LTQ Orbitrap Discovery mass spectrometer coupled to an Agilent 1200 series HPLC. Labeled peptide samples were pressure loaded onto 250 mm fused silica desalting column packed with 4 cm of Aqua C18 reverse phase resin (Phenomenex). Peptides were eluted onto a 100-mm fused silica biphasic column packed with 10 cm C18 resin and 4 cm Partisphere strong cation exchange resin (SCX, Whatman), using a five-step multidimensional LC-MS protocol (MudPIT). Each of the five steps used a salt push (0%, 50%, 80%, 100%, and 100%), followed by a gradient of buffer B in Buffer A (Buffer A: 95% water, 5% acetonitrile, 0.1% formic acid; Buffer B: 20% water, 80% acetonitrile,

0.1% formic acid) as outlined previously.[9] The flow-rate through the column was ~0.25μL/min, with a spray voltage of 2.75 kV. One full MS1 scan (400-1800 MW) was followed by 8 data dependent scans of the nth most intense ion. Dynamic exclusion was enabled. The tandem MS data, generated from the 5 MudPIT runs, was analyzed by the SEQUEST algorithm.[7] A static (+57.0215 m/z, iodoacetamide alkylation) modification on cysteine residues was specified. The precursor-ion mass tolerance was set at 50 ppm while the fragment-ion mass tolerance was set to 0 (default setting). Data was searched against an *E. coli* reverse-concatenated non-redundant FASTA database containing Uniprot identifiers. MS datasets were independently searched with light and heavy parameter files; for these searches, differential modifications on cysteine for either the IAL (+306.1481) or IAH (+312.1682) modification were used. MS2 spectra matches were assembled into protein identifications and filtered using DTASelect2.0,[8] to generate a list of protein hits with a peptide false-discovery rate of <5%. With the –trypstat and –modstat options applied, peptides were restricted to fully tryptic (-y 2) with a found modification (-m 0) and a delta-CN score greater than 0.06 (-d 0.06). Single peptides per locus were also allowed (-p 1) as were redundant peptide identifications from multiple proteins, but the database contained only a single consensus splice variant for each protein. Quantification of peptide L/H ratios were calculated using the cimage quantification package described previously.[9] Annotation of protein subcellular localization as well as cysteine function and conservation were generated from the Uniprot Protein Knowledgebase (UniProtKB) as described previously.[12]

**Chemoproteomic labeling of E. coli cysteine residues for reactivity profiling.** For reactivity profiling of *E. coli* cysteine residues (**Supplemental Tables 7**), 2 mg of iron-depleted *E. coli* (K-12) lysate were labeled with either 100 μM IA-alkyne Light (IAL – 10 μL, 10 mM stock in DMSO) or 10 μM IA-alkyne Heavy (IAH – 10 μL, 1 mM stock in DMSO) for 1 h at room temperature. Probe labeled proteins were then conjugated to chemically-cleavable diazobenzene biotin-azide (azo-tag) by Cu(I)-catalyzed [3 + 2] cycloaddition. The samples were then carried through the same sample preparation as described previously for isoTOP-ABPP analysis and analyzed as described by LC/LC-MS/MS on a Thermo LTQ Orbitrap Discovery mass spectrometer coupled to an Agilent 1200 series HPLC.

**Purification and characterization of recombinant TrhP**. Cell pellets were resuspended in 5 mL TrhP lysis buffer (PBS, 5 mM imidazole, 1 mM DTT), and lysed by sonication (3 x10 pulses – 90%) on ice. Cell lysates were centrifuged at 10,000 g for 10 min at 4 ºC to remove unlysed cell debris. The resulting supernatant was loaded onto 1 mL of Ni-NTA that had been equilibrated with lysis buffer. The Ni-NTA was washed with 5-bed volumes wash buffer (PBS, 20 mM imidazole, 1 mM DTT). Recombinant AcnA was eluted from the Ni-NTA column with 2-bed volumes of elution buffer (PBS, 250 mM imidazole, 1 mM DTT). The protein was desalted over a pre-packed PD-10 column equilibrated with PBS buffer and concentrated using an Amicon-Ultra 10 kDa centrifugal filter. Protein concentration was determined by Bradford assay, and the presence or absence of an Fe-S cluster was confirmed by UV-visible absorption spectroscopy on a ThermoScientific Nanodrop 2000c spectrophotometer. Recombinant TrhP was diluted to 1 μg $mL^{-1}$ in 5% of TraceMetal Grade Nitric Acid in Milli-Q double distilled water. Iron content was measured on an Agilent 5100 ICP-OES. Samples were prepared in triplicate and the relative iron concentration of each cell pellet sample was calculated from a calibration curve of external standards (iron – 0 to 56 mg $l^{-1}$) according to peak height.

**Purification and characterization of recombinant DppD**. Recombinant DppD was purified and characterized in the exact same manner as TrhP, with the exception that 5% glycerol and 1% Triton-X were added to the lysis, wash, and elution buffer during protein purification.
